# Droplet-based single cell RNA sequencing of bacteria identifies known and previously unseen cellular states

**DOI:** 10.1101/2021.03.10.434868

**Authors:** Ryan McNulty, Duluxan Sritharan, Shichen Liu, Sahand Hormoz, Adam Z. Rosenthal

## Abstract

Clonal bacterial populations rely on transcriptional variation to differentiate into specialized cell states that increase the community’s fitness. Such heterogeneous gene expression is implicated in many fundamental microbial processes including sporulation, cell communication, detoxification, substrate utilization, competence, biofilm formation, motility, pathogenicity, and antibiotic resistance^1^. To identify these specialized cell states and determine the processes by which they develop, we need to study isogenic bacterial populations at the single cell level^2,3^. Here, we develop a method that uses DNA probes and leverages an existing commercial microfluidic platform (10X Chromium) to conduct bacterial single cell RNA sequencing. We sequenced the transcriptome of over 15,000 individual bacterial cells, detecting on average 365 transcripts mapping to 265 genes per cell in *B. subtilis* and 329 transcripts mapping to 149 genes per cell in *E. coli*. Our findings correctly identify known cell states and uncover previously unreported cell states. Interestingly, we find that some metabolic pathways segregate into distinct subpopulations across different bacteria and growth conditions, suggesting that some cellular processes may be more prone to differentiation than others. Our high throughput, highly resolved single cell transcriptomic platform can be broadly used for understanding heterogeneity in microbial populations.

## Introduction

In recent years, single cell RNA sequencing (scRNA-seq) has enabled the agnostic assessment of transcriptional heterogeneity in eukaryotic systems. Since the introduction of microfluidic techniques which allow encapsulation of cells and tagging of poly-adenylated transcripts in small droplets, scRNA-seq studies of mammalian populations have grown exponentially. However, tools to comprehensively study differential gene expression in bacterial populations remain severely limited due to several significant technical challenges. First, total mRNA abundance in bacteria is two orders of magnitude lower than that of eukaryotes, with a single bacterial cell containing approximately 10^3^ - 10^4^ transcripts during exponential growth^4^. Second, transcriptional turnover is much faster in bacteria with mRNA half-life often on the scale of minutes, compared with hours for eukaryotic transcripts^4^. Third, bacterial transcripts do not intrinsically include a 3’ poly-adenosine tail and, therefore, mRNA cannot easily be tagged and selectively enriched against rRNA, which makes up >95% of the bacterial transcriptome^4^. Lastly, accessing mRNA requires cell permeabilization which is markedly more difficult in bacteria due to the diversity in membrane structure and variability of the peptidoglycan layer. We reasoned that a new method combining the advantages of microfluidic single cell barcoding in droplets with the ability to tag transcripts using in-situ hybridization of oligonucleotide probes could overcome these challenges. Here, we present a method for prokaryotic single cell RNA sequencing which utilizes a commercial, benchtop microfluidic device (Chromium Controller from 10X Genomics) and custom ssDNA probe libraries to resolve the mRNA profile of thousands of Gram positive (*Bacillus subtilis* 168) and Gram negative (*Escherichia coli* MG1655) bacterial cells.

## Method Development

### Probe Design & Library Generation

To leverage existing microfluidic single cell sequencing platforms, we devised a method relying on tagging of individual transcripts with DNA probes. This approach requires the generation of a large oligonucleotide library that is complementary to all protein coding sequences within a genome (Figure 1). Multiple DNA regions of 50 bp were chosen from each ORF based on uniqueness as determined by UPS2 software^5^ or based on previously published oligonucleotide arrays^6,7^. These sequences then served as the hybridization regions of ssDNA probes which were designed to target mRNA by sequence complementarity. ssDNA probes also contained a 5’ PCR handle for library generation, a Unique Molecular Identifier (UMI), and a 3’ poly adenosine tail (A_30_) for retrofitting prokaryotic transcripts to the Chromium Single Cell 3’ system (Figure 1a). Multiple probes (complementary to different regions) were designed for each gene to enhance transcript capture efficiency and decrease noise caused by poor hybridization and/or insufficient amplification of any given probe. Complete species libraries contained 29,765 probes for *B subtilis* and 21,527 probes for *E. coli*, and targeted 2,959 and 4,181 genes, respectively. Libraries were ordered at sub-femtomole quantities from Twist Biosciences (a one-time cost of 5¢ - 15¢ per probe) and amplified by rolling circle amplification^8^ to obtain a sufficient concentration (0.25 mg = 10.25 nM/library or approximately 0.35 pM of each probe) for scRNA-seq experiments (Figure1b, Methods, Supplementary Tables S1 and S2, Supplementary Figure S1). Probe libraries were completed by addition of randomized 12 bp UMI sequences and a poly-A tail and purified by PAGE. Completed libraries had a uniform coverage of probes (Figure 1e). The rolling circle amplification approach permits re-amplifying probe sets for unlimited subsequent experiments without the need to re-order probes, at an upfront cost of under 1¢ per cell (Supplementary Table S3).

**Figure 1.**
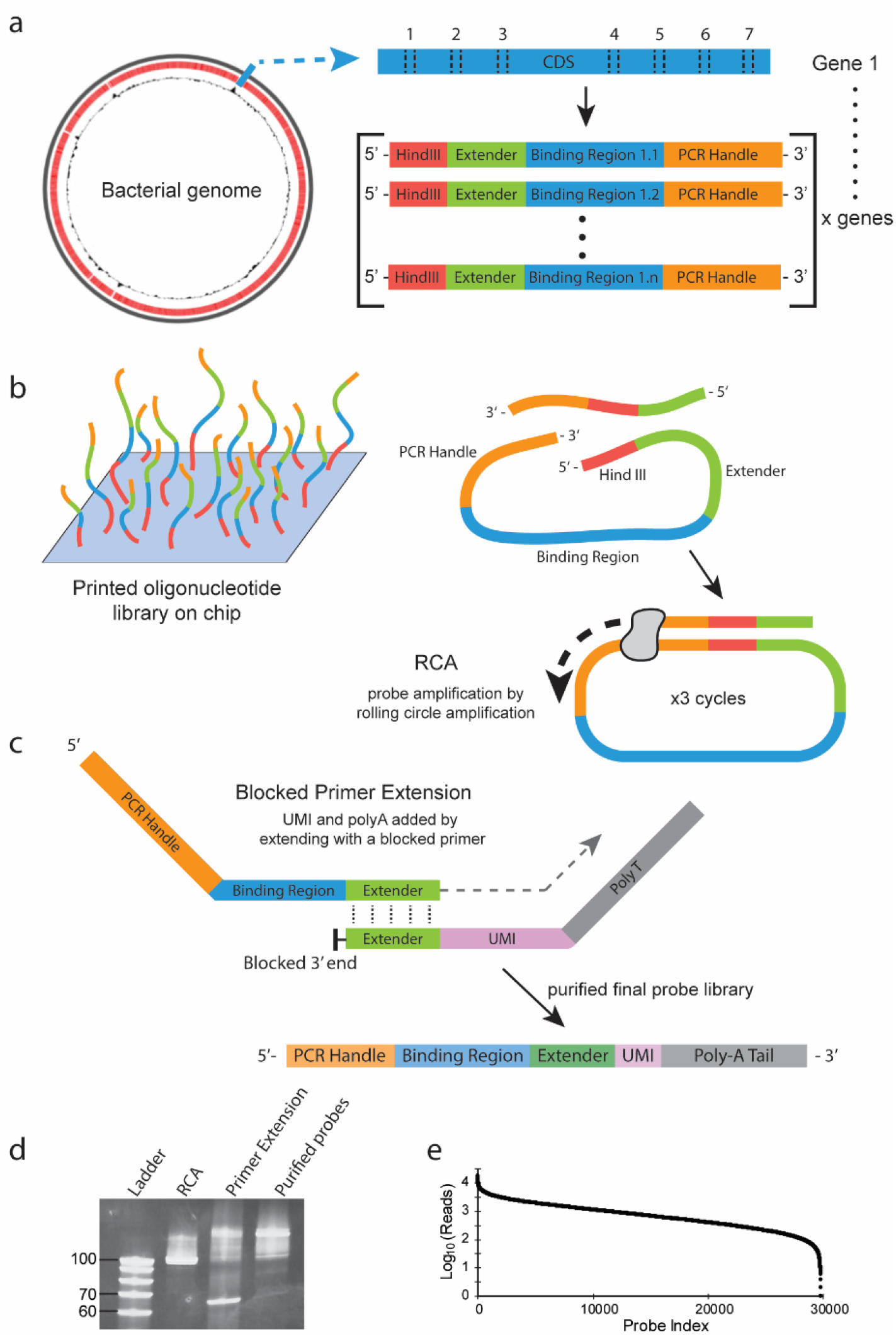
Genome wide scRNA-seq probe design and synthesis. **a**. A genome for the organism of interest is used to design a library of probes that are complementary to unique regions within each gene’s coding sequence (CDS). Complementary ssDNA oligonucleotides are amended by addition of an extender sequence at the 5’ end and 3’ PCR handle. Sequences are ordered as an oligopool **b**. Probe sequences are inverted in-silico and restriction sites and handles are added for circle-to-circle amplification **c**. The large, inverse oligonucleotide library is commercially-synthesized and pooled. Ordered oligo-pools are amplified in a 3-step “rolling circle” reaction with Phi29 polymerase to produce the final, correctly oriented proto-probe library. **d**. A random 12mer sequence (UMI) and a 30-base polyA tail is added by primer extension with a blocked primer that is complementary to the 3’ extender sequence. Single stranded DNA containing a universal PCR handle, UMI, complementary transcript region, and poly-A tail (listed 5’ to 3’) is purified. **e**. Finalized probe libraries are free of amplification primers and well distributed.

Before microfluidic encapsulation, bacteria were fixed in 1% paraformaldehyde and permeabilized (Methods). Permeabilized bacteria were incubated with their corresponding DNA probe library. Non-hybridized probes were washed away (Figure 2a, Methods). Next, the bacteria were run through a 10X controller, where the DNA probes were captured and barcoded in a manner analogous to the barcoding of the transcriptome of mammalian cells (Figure 2 b-d). The resulting libraries were sequenced, preprocessed with custom scripts (https://gitlab.com/hormozlab/bacteria_scrnaseq), and analyzed with the standard CellRanger pipeline and the Seurat analysis package^9,10^ (see Methods).

**Figure 2.**
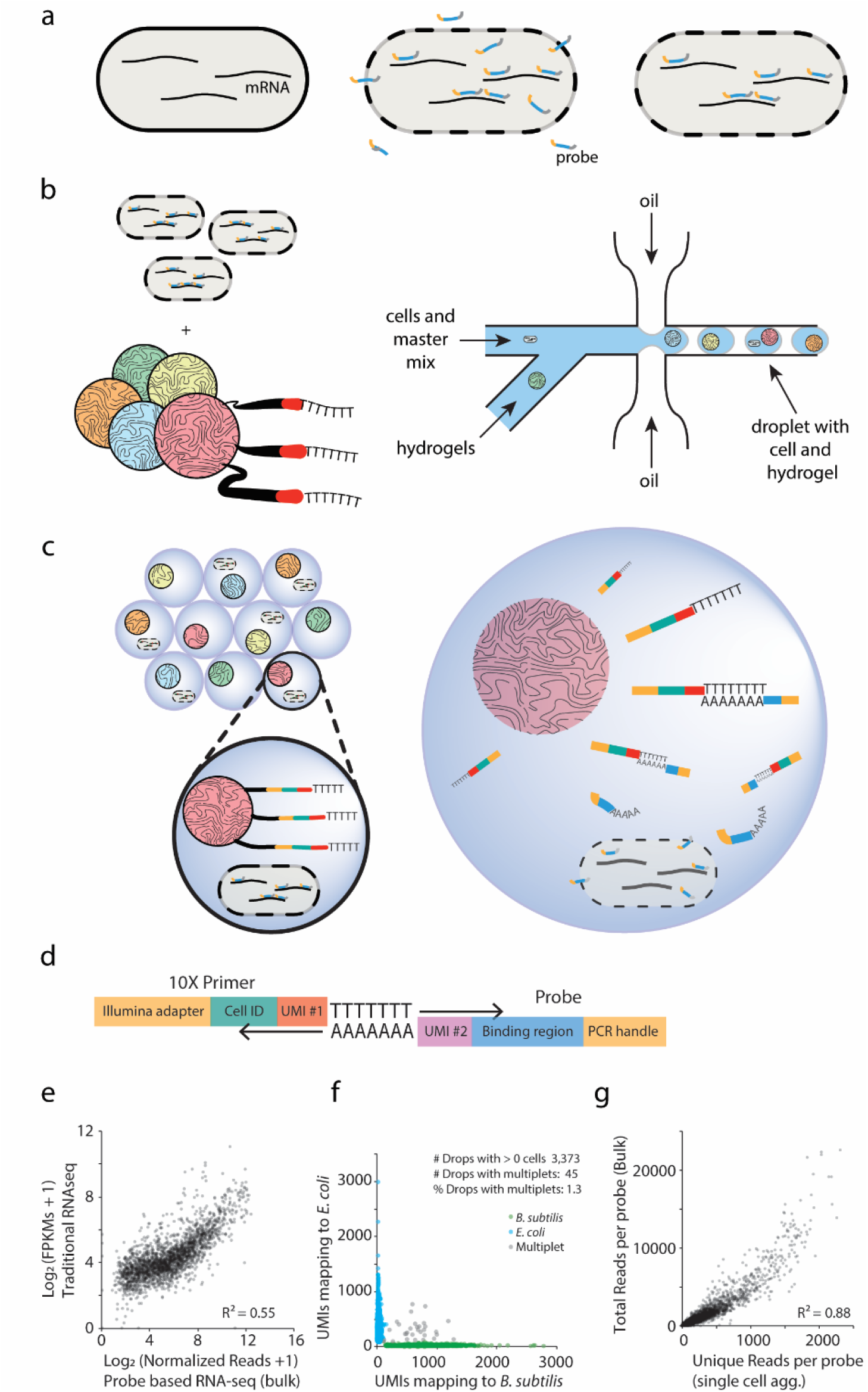
Microfluidic probe-based scRNA-seq method and validation. **a**. Cells are fixed and permeabilized to allow penetration of thousands of unique, genome-specific oligonucleotide probes. Hybridized probes “retrofit” transcripts with a poly-A tail & UMI while unhybridized probes are washed away. **b**. Cells are then flowed through a commercial microfluidic device which encapsulates single cells into droplets (shown in **c**.) containing barcoded primers conjugated to a hydrogel microsphere and PCR reagents. **d**. Barcoded cDNA is generated from the mRNA:probe hybridized complex via in-droplet PCR or RT reaction. Droplets are then broken, the pooled cDNA amplified further before sequencing. Single cell transcriptomes are resolved, clustered, and visualized. **e**. Transcriptomic estimation by probe hybridization in bulk samples is correlated to transcriptomic measurement of bulk samples by traditional RNA-seq **f**. Species mixture (“barnyard”) plot demonstrates that single cells of different bacterial species can be resolved by barcode after microfluidic encapsulation. **g**. Aggregated probe-based signal from thousands of single cells correlates with the average probe-based signal of the bulk population.

### Microfluidic RNA-seq validation in bulk and single cell experiments

To determine if our probe library can report on the transcriptional state of cells, we split a culture of *B. subtili*s cells in late exponential state into two aliquots of approximately 10^8^ cells each. One aliquot was processed using a traditional RNA-seq protocol (Methods) while the second sample was fixed in formaldehyde for *in situ* hybridization with a probe set designed against the *Bacillus subtilis* genome. After incubation and washing, the probes that remained bound were amplified by PCR and processed into Illumina libraries. The output was compared to the traditional RNA-seq library. The hybridization and wash conditions at 50°C gave results that are similar to traditional RNA-seq (Figure 2e; R^2^ = 0.55). This value is comparable to the correlation observed when comparing RNA-seq to microarray experiments (R^2^ = 0.57-0.65)^11,12^.

Next, we tested if single cell transcriptomes can be captured by encapsulating individual bacteria using the 10X Chromium Controller after fixation and *in situ* probe hybridization. *E. coli* cells and *B. subtilis* cells were independently pre-treated with probes corresponding to their respective genomes. The prepared bacterial samples were subsequently mixed and loaded onto a microfluidic chip along with a custom PCR master mix containing a DNA primer designed to amplify the back-end PCR handle built into the probes (Figure 1, Methods). This custom mix replaced the standard cDNA reagents supplied by 10X Genomics as the aqueous phase within the droplets. Sequenced libraries from this experiment demonstrate that the microfluidic platform is able to successfully segregate and barcode individual microbial cells (Figure 2f). We observed a “doublet” rate of 1.3% for 3,373 captured cells, consistent with the expected rate of successful single cell encapsulation based on the specifications of the 10X microfluidic system (reported as an expected 2.3% multiplet events per 3,000 captured cells^10^).

To assess whether the signal from individually encapsulated cells provides an unbiased readout of the transcriptomic state of the population, we compared the output produced by capturing probes from a bulk sample of cells (≅10^8^ cells) to the signal obtained by summing the probe counts over thousands of individually tagged cells (aggregated UMI counts). The number of reads across all probes between the two samples were highly correlated (R^2^ = 0.88, Figure 2g), confirming that the signal from single cells provides an unbiased representation of transcriptional states.

## Results

### Bacterial scRNA-seq identifies known and previously hidden heterogeneity in *B. subtilis*

We performed scRNA-seq on *B. subtilis* grown to late exponential phase in M9 minimal media supplemented with malate. In total, 2,784 cells were captured as determined by unique barcodes with 550±340 (mean±SD, median=470) reads per cell. We detected a median of 325 mRNA transcripts per cell (approximately 10-15% of the total mRNA pool^13^) corresponding to a median of 241 genes per cell. UMIs per probe ranged from 0-45 for any given gene in any given cell. The most highly detected probes targeted genes encoding ribosomal proteins (rpsG, rpsH, rpsR, rpsM, rpsS, rplL, rplD, rplO, rplV), translation elongation machinery (fusA, tufA, map), and transcriptional machinery (rpoA).

Next, we resolved distinct cellular states by dimensional reduction of single cell expression vectors followed by graph-based clustering and analysis of differential gene expression (DGE) using the Seurat package^9^. For *B. subtilis* in M9 minimal media in late log growth, we resolved four major transcriptomic signatures comprised of ten cell clusters that capture more subtle variations in gene expression (Figure 3 a-b, Supplementary Figure S2, Supplementary Table S6). Algorithmic grouping of cell expression profiles was robust to the clustering parameters (see Methods). As expected for minimal media cultures in late log, we observed a subpopulation of cells (clusters 6 and 8, approximately 10% of all cells) in the genetically competent state, the physiological state in which bacteria can uptake extracellular DNA and incorporate it into their genome. Cells in these two clusters differentially overexpress the competence master-regulator comK (logFC = 2.37, adjusted p-value = 4E-179) as well as multiple genes within and downstream of competence operons (comC, comEA, comEC, comFA, comFB, comFC, comGA, comGC, comGD, comGE, comGG, recA, coiA, dprA, sbB, nucA, nin, clpC, recA). The fraction of competent cells in the population was measured by fluorescence microscopy using a fluorescent promoter-reporter of comG (Methods, Figure 3c left panel, Supplementary Figure S12a) and determined to be approximately 10.4% (IQR outlier detection - Methods) - similar to the fraction of the population comprising clusters 6 and 8 (9.3%, Supplementary Table S5). In total, we found that 45 of the 50 comK-regulon genes that were probed were differentially overexpressed in clusters 6 and 8 (adjusted p-values < 0.05; Figure 3d, Supplementary Table S6). Our results agree with numerous studies identifying the competence regulon in *B. subtilis*^14,15^, and the percentage of cells displaying natural competence falls within the range (3-10%) of previous observations in similar media^16,17^.

**Figure 3.**
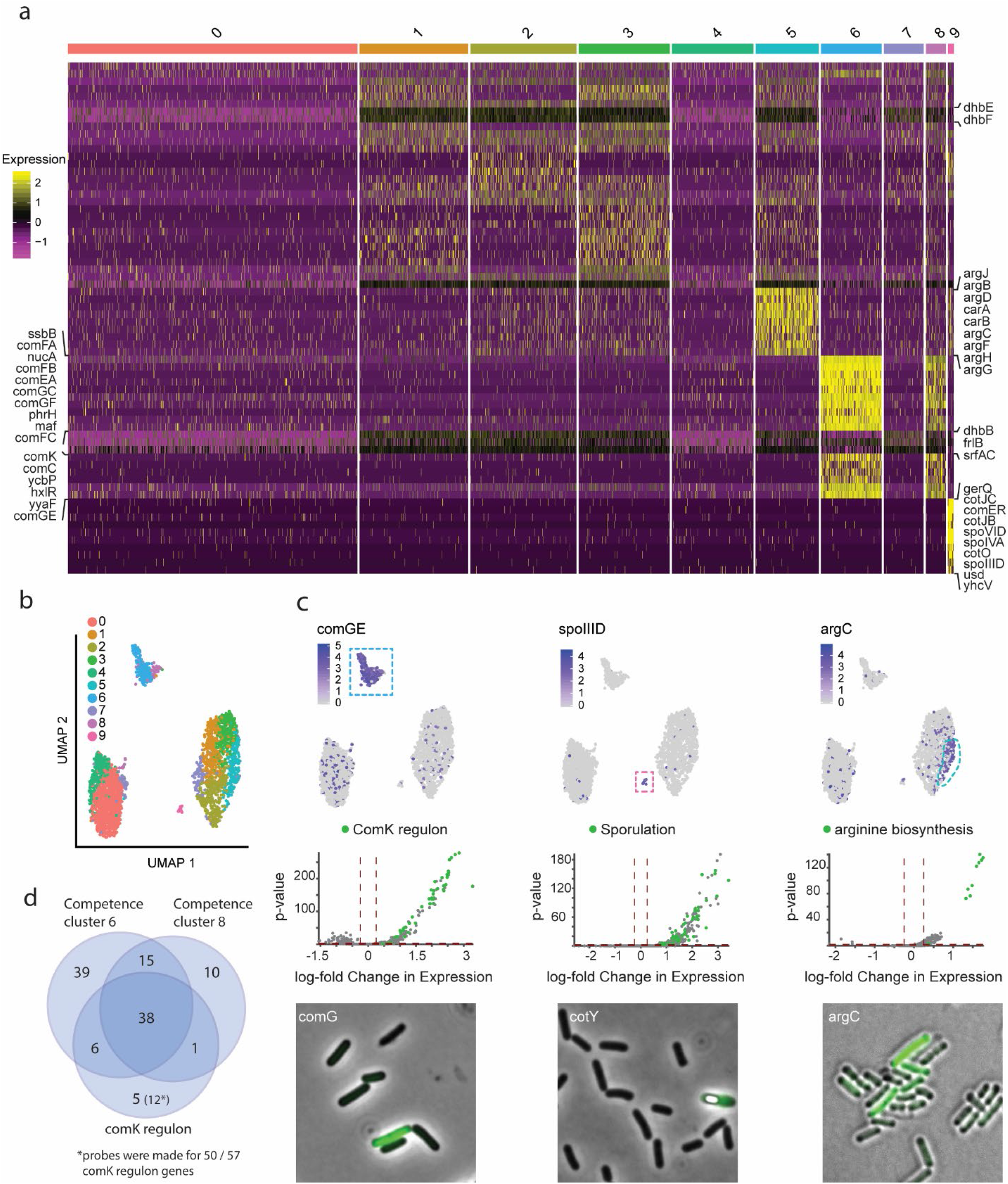
scRNA-seq analysis reveals known and novel states in *B subtilis*. **a**. Heatmap of marker gene expression (z-score of log-transformed values) from 2,784 individual *B. subtilis* cells organized into 10 clusters. **b**. UMAP 2-dimensional representation of the 10 cell clusters reveals 4 highly distinct transcriptomic signatures **c**. Single cell expression of key marker genes for competence (comGE; clusters 6 and 8), sporulation (spoIIID, cluster 9) and arginine synthesis (argC, cluster 5) are highlighted on the UMAP (top panels, left to right). Volcano plots of genes expressed in at least 25% of cells in the corresponding clusters, with genes from the respective processes highlighted in green (middle panels). The lower panels show the presence of each heterogeneous marker in the population as confirmed by fluorescent promoter-reporter constructs (P_*comG*_-YFP, P_*cotY*_-YFP, P_*argC*_-YFP, respectively). A phase bright spore can be seen in the cotY-expressing cell. **d**. 90% of all comK regulon genes probed (45/50) were significantly differentially upregulated in cell clusters 6 and 8 (Bonferroni corrected P-value ≤ 0.05 for each gene)

Unexpectedly for these growth conditions, we observed a small subpopulation of cells within the sample (cluster 9, 19 cells, 0.7% of the population) with a transcriptomic signature indicative of sporulation. In these cells, we observed significant upregulation of transcripts corresponding to sigma factors associated with sporulation, sigF (logFC = 1.2, adjusted p-value = 4E-41) and sigG (logFC = 2.2, adjusted p-value = 2E-63), as well as genes in the ger, cot, and spoIVF operons within the cluster. Gene set enrichment analysis (GSEA) on all significantly upregulated genes in cluster 9 using ontological classes from the Gene Ontology Consortium ^18–20^ reveals a 4.3-fold enrichment of genes involved in sporulation (Fisher’s exact test, FDR = 1E-16) as well as spore germination (fold enrichment = 8.1, FDR = 1.7E-03). This finding highlights the ability of probe-based scRNA-seq to detect rare and unexpected cell states in microbial cultures. To confirm this subpopulation, we created a promoter reporter for cotY, a spore coat gene expressed in the sporulating mother cell. Fluorescence observation identifies the presence of cotY expression and spores in a similar percentage of cells as recovered by scRNA-seq (0.5-1%) (Figure 3c middle panel, Supplementary Figure S12d).

The largest group of *B. subtilis* cells (comprising clusters 1, 2, 3, 5, and part of 7) accounted for ∼45% of the population and was characterized primarily by the upregulation of the dhb operon (logFCs > 0.25, adjusted p-values < 2E-05). Genes dhbA, dhbB, dhbC, dhbE, and dhbF encode the 5 enzymes implicated in the biosynthesis of bacillibactin – a catecholic siderophore which is produced and secreted in response to iron deprivation inside the cell. Relatedly, yusV, an ABC transporter of bacillibactin, was also found to be upregulated in these cells. Cells within cluster 5 are distinguished from the rest of the subpopulation by expression of genes related to arginine biosynthesis via ornithine. 10 of 14 genes associated with the cellular process were significantly upregulated in this cluster alone (gene set fold enrichment = 19.35, FDR = 3E-08). A fluorescent promoter reporter for argC confirms heterogeneous expression of arginine genes with approximately 2.1% of cells in the high-argC expressing tail of the distribution - compared to 7.2% of cells in this state as determined by scRNA-seq (Figure 3c right panel, Supplementary Figure S12b, Methods, Supplementary Table S5).

Finally, to see if microfluidic probe-based methods are robust with different DNA-labeling chemistries, we repeated scRNA-seq on *B. subtilis*, replacing in-droplet PCR with a probe barcoding step utilizing a reverse transcriptase enzyme in accordance with the 10X Chromium Single Cell 3’ standard protocol for eukaryotic scRNA-seq. While resolution decreased (median of 89 transcripts and 86 unique genes per cell), we were still able to resolve distinct biological states including sporulating and competent cell clusters (Supplementary Figure S3, Supplementary Table S6). Taken together, our platform for single cell sequencing of individual bacterium correctly recapitulates the known cellular states of *B. subtilis* at the expected population fractions and identifies previously unknown cellular states.

### Gene expression heterogeneity in clonal populations of *E. coli*

With a validated method for identifying cellular states in *Bacillus subtilis*, we turned our attention to characterizing transcriptional heterogeneity in *Escherichia coli. E. coli* MG1655 was first grown in M9 minimal media, fixed at an OD_600_=0.5 (mid log), and subjected to probe-based scRNA-seq. We resolved 3,315 cells by unique single cell barcodes, detecting a median of 263 transcripts per cell representing a median of 165 distinct genes. Cells were partitioned into nine groups by graph-based clustering of their gene expression profiles (Figure 4ac, Supplementary Figure S4, Supplementary Table S6). The lack of clearly distinguishable transcriptomic signatures after dimensionality reduction using UMAP suggests a cell population that exhibits more uniform gene expression profiles compared with *Bacillus subtilis*. However, as with *B. subtilis*, we observe that certain clusters can be assigned to specific biological processes by DGE and that cluster determination is robust to the choice of clustering parameters (Methods). Cells in cluster 8 (141 cells, 4.3% of the population) exhibit significant upregulation of fimI, fimC, and fimA (logFC > 2.4, adjusted p-value < 1E-166). The downstream genes of the fim operon encode for components of type 1 pili and are known to be expressed heterogeneously in cells upon inversion of the DNA element containing the fimA promoter (“phase on” fim switch) ^21,22^. Type 1 pili are associated with biofilm formation and pathogenicity due to the adhesive properties of the fimbriae ^23^.

**Figure 4.**
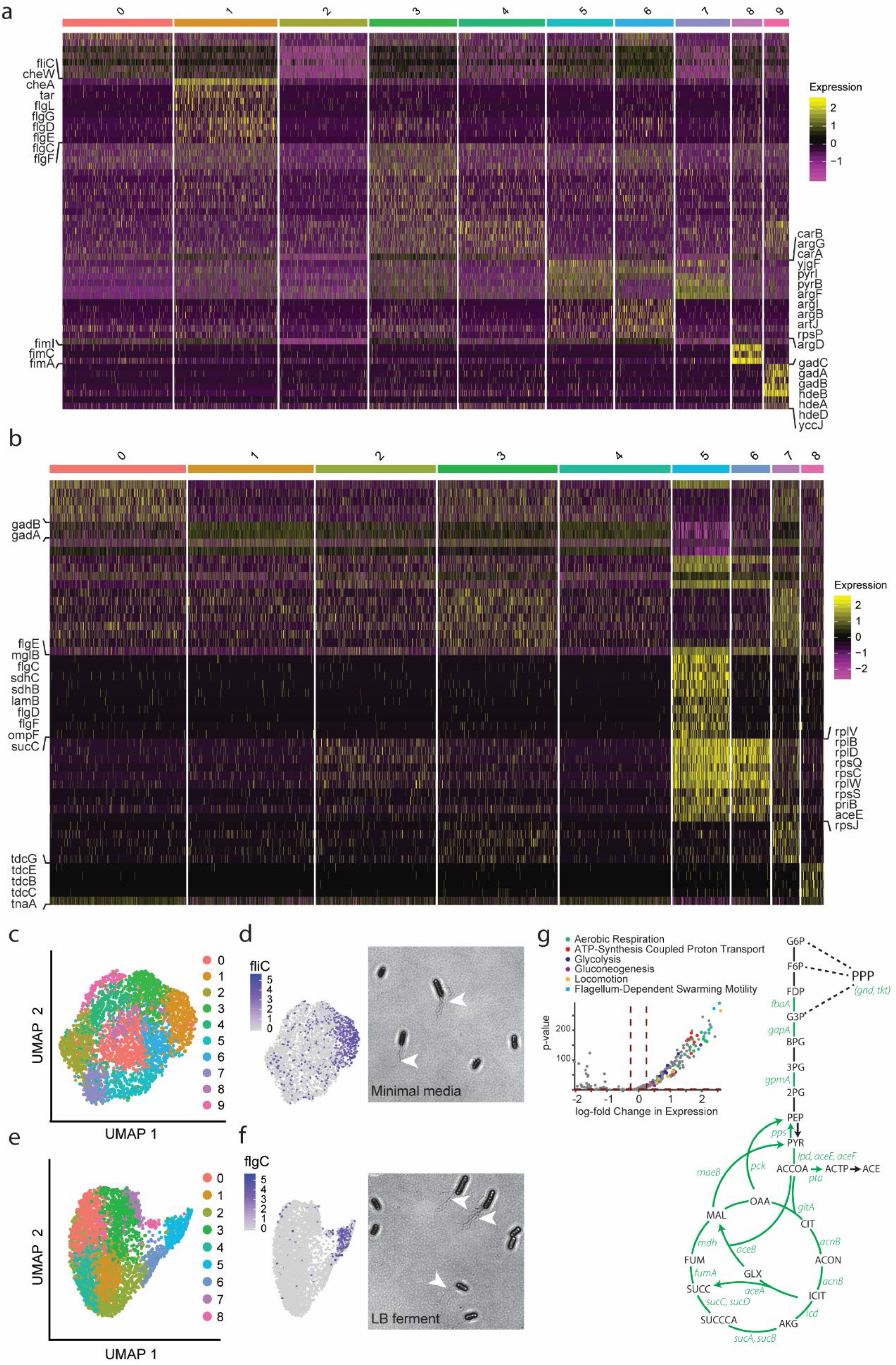
Heterogeneous gene expression in *E coli* grown in minimal medium and anaerobically. **a**. Heatmap of marker gene expression (z-score of log-transformed values) from 3,315 individual *E. coli* cells grown aerobically in minimal M9 media organized into 10 clusters. **b**. Heatmap of gene expression from 3,763 individual *E. coli* cells grown in anaerobic LB media organized into 9 clusters. **c**. UMAP 2-dimensional representation of the 10 cell clusters from aerobic M9 culture conditions. **d**. Flagellar components are heterogeneously expressed in aerobic M9 media - a key flagellar gene (flagellin - fliC) is preferentially expressed by cells in cluster 1. The heterogeneous presence of flagella in the population is confirmed by flagellar staining. **e**. UMAP 2-dimensional representation of the 9 cell clusters from *E coli* fermenting LB media culture conditions. **f**. Flagellar components are heterogeneously expressed in fermentation conditions - a key flagellar gene (flagellar basal-body rod protein - flgC) is preferentially expressed by cells in cluster 5. The heterogeneous presence of flagella in the population is confirmed by flagellar staining. **g**. Many aerobic respiration genes and other metabolic and flagellar genes are overexpressed in cluster 5. Volcano plots of genes over-expressed in at least 25% of cells in cluster 5 from these processes are highlighted in colors as labeled. Green labeled genes, involved in central-carbon metabolism, are also highlighted as green arrows in the metabolic map.

Relatedly, cells in cluster 1 (482 cells, 14.5% of population) uniquely upregulate genes implicated in cell motility including those encoding chemotaxis signaling proteins (cheA, cheW, tar; gene set fold enrichment > 100, FDR = 3.1E-02) and structural flagella components (fliC, flgL, flgG, flgD, flgE, flgC, flgF; gene set enrichment = 86.08, FDR = 2.6E-06). This list includes genes regulated by class 1, 2, and 3 flagella promoters; three distinct regulons which control the sequential expression of flagella genes including the master regulator flhDC, components of the membrane-associated basal body and chemotaxis proteins, respectively^44^. This model of assembly has been described as “just in time” gene expression that allows *E. coli* to discretize the energetically demanding biosynthesis process into checkpoint-controlled subprocesses. The fact that we observe upregulation of genes from multiple classes within the cluster, often within the same individual cells, indicates a more overlapping transcriptomic state, an observation that aligns with the fact that class 2 controlled fliA controls transcription of class 3 promoters upon completion of the basal body and export of class 2 flgM (fliA inhibitor)^44^. To confirm this motile subpopulation of cells, we stained cells grown under the same conditions (Remel flagella stain) and visualized by phase microscopy. As shown in Figure 4d, we see assembled flagella emanating from a fraction of cells within the population.

Cells within clusters 5, 6, and 7 demonstrate differential usage of carbamoyl phosphate within the culture. Clusters 5 and 7 upregulate carAB, which encode the subunits of carbamoyl phosphate synthetase, converting bicarbonate to carbamoyl phosphate as the first step in the de novo biosynthesis of both uridine-5’-monophosphate (UMP) and arginine. Both clusters, arranged adjacently on the UMAP, also upregulate pyrI, pyrB & pyrC, three enzymes involved in the first two steps of UMP biosynthesis, implying a commitment of the cells towards pyrimidine biosynthesis. Alternatively, cells in cluster 6 shuttle carbamoyl phosphate into the arginine biosynthesis pathway as implied by the upregulation of argF, argI, argB, argG, argD (logFCs > 0.55, adjusted p-values < 1E-20). GSEA reinforces this observation, finding a significant enrichment of genes implicated in arginine biosynthesis via ornithine in these cells (gene set fold enrichment = 26.77, FDR = 2.02E-03).

Next, *E. coli* was grown in rich, LB media under anaerobic conditions. In the absence of exogenous electron acceptors permitting cellular respiration (e.g. O2, nitrite, nitrate, DMSO), *E. coli* performs “mixed acid” fermentation to sustain ATP generation by glycolysis, shuttling electrons into a variety of end-products (acetate, ethanol, succinate, formate) in order to replenish the pool of NAD+. The amount of each end-product produced by *E. coli* varies based on the redox state of the cell - an adaptability which allows *E. coli* to grow on a wide range of substrates under anoxic conditions and varying pH^24^. We performed scRNA-seq to elucidate whether mixed acid fermentation can be, in part, attributed to the separation of branched fermentative pathways into distinct subpopulations of cells. Graph-based clustering of gene expression vectors (n = 3,763 resolved cells) resulted in 8 cell clusters (Figure 4b,d, Supplementary Figure S5, Supplementary Table S6). Within the population, we observe the most highly abundant transcripts (top 5% by aggregated UMIs) corresponding to genes involved in anaerobic ATP generation including glycolysis (fbaB, mdh, aceE, fbaA, pgi, lpd, eno, gapA, pgk, gene set fold enrichment = 9.5 compared with a null model where the top 5% most abundant genes were chosen randomly, FDR = 2.0E-4), anaerobic respiration (dcuA, acnA, hyaC, glpB, frdC, mdh, frdB, hyaA, hyaB, narH, glpD, glpA, narG, hybO, glpC, fold enrichment = 4.9, FDR 1.22-04), and mixed acid fermentation (hyaC, frdC, mdh, frdB, hyaA, hyaB, gldA, fold enrichment = 8.3, FDR = 3.7E-03) in addition to genes related to ribosome biogenesis, transcription, and translation. Enrichment of these pathways within the most highly expressed genes at the population level is consistent with an anaerobic environment.

We then analyzed the transcriptional states of cells in each cluster. Clusters 1 and 4 appear to represent a subpopulation of cells responding to over-acidification of the cytoplasm. Both groups significantly upregulate the gad operon (gadABC) as well as dps (logFCs > 0.34, adjusted p-values < 1E-32). In addition, acid stress chaperone proteins hdeA and hdeB are overexpressed in cluster 4 (logFCs > 0.25, adjusted p-value < 1E-02). Acid stress is an expected consequence of fermentation as acidic fermentation byproducts (e.g. acetate, lactate, formate) accumulate. Cells in cluster 5 (283 cells, 7.5% of total) had 157 genes significantly upregulated (adjusted p-values <0.05) compared with the rest of the population. Surprisingly, these genes belong to pathways involved in aerobic respiration (aceAB, sucABCD, nuoBMN, cyoABC, cydAB, sdhABC, fumA, gltA, acnB, icd, mdh; gene set fold enrichment = 11, FDR = 2.32E-13), ATP synthesis coupled proton transport (atpABCDEHG; gene set fold enrichment = 24.47, FDR = 1.45E-05), glycolysis (mdh, gpmA, aceE, fbaA, lpd, aceF, gapA; gene set fold enrichment = 10.24, FDR = 7.33E-04), and gluconeogenesis (gpmA, fbaA, sdaB, pck, pps; gene set fold enrichment = 9.92, FDR = 1.03E-02). The overexpression of these genes, many of which are involved in the TCA cycle (Figure 4g), may be an indication of oxygen contamination within the culture, resulting in a stark difference in transcriptional signatures and, consequently, cell behavior within the isogenic population. In addition, cells in cluster 5 also appear to be motile as genes pertaining to flagella structure (flgCDEFG; fold enrichment = 9.26, FDR = 1.28E-02) and locomotion (fliCD, tar, mglB, cheAW, flgK; fold enrichment = 5.16, FDR = 4.08E-04) were upregulated & overrepresented in the subpopulation. Heterogeneous expression of flagella was confirmed microscopically using a remel stain (Figure 4f). Since cultures were maintained in capped tubes outside of an anaerobic chamber, is it possible air was introduced into the culture through an imperfect seal. Thus, we hypothesize that this induced cell population activated motility genes to gather at the liquid/air interface. Consistent with the idea of such an oxygen-gradient organization, a recent report used a high-throughput fluorescent *in situ* hybridization (FISH) analysis of approximately 100 genes to demonstrate the role of oxygen gradients in spatial partitioning within biofilms^25^.

Finally, cluster 8 (111 cells, 3% of total), which projects between the majority of cells and cluster 5 on the UMAP, is characterized by the strong upregulation of the tdc operon (tdcBCEG; logFC > 3.0, adjusted p-values < 1E-100). Under anaerobic conditions, if energy is low inside the cell, *E. coli* generates ATP by conversion of L-threonine to propionate. The tdc operon has been shown to be strongly induced by anaerobic conditions as well as catabolically repressed (CRP-controlled), suggesting these cells are sensing an anoxic environment while also deprived of glucose^26^. The tdc operon encodes proteins for the transport and degradation of L-threonine (and L-serine by analogous intermediates) for the purpose of ATP generation (1:1 conversion). While the operon is induced in the absence of oxygen, it has been shown to not be directly regulated by common oxygen-sensing systems inside the cell such as transcriptional regulators FNR and ArcA but rather by tcdA and tcdR (transcriptional activators) ^27^. To the best of our knowledge, this is the first time threonine metabolism has been shown to be cellularly segregated in a microbial culture. Taken together, these findings demonstrate how single cell transcriptomics can inform on physiological and metabolic organization in complex microbial environments.

## Discussion

To agnostically characterize the collection of cell states within a microbial population, we developed an affordable (standard commercial 10X run costs plus ≅ 1¢ per cell in probes) bacterial scRNA-seq technique that demonstrates high throughput and sensitivity. Our method, which is based on combining microfluidic droplet partitioning of single cells with *in situ* hybridization of DNA probes, overcomes several difficulties associated with obtaining quality single cell mRNA signal from bacteria. Using bacterial scRNA-seq, we identify known cellular states including genetic competence and sporulation in *B. subtilis* and fimbriae production in *E coli*. In addition to these well characterized physiologies, we uncovered several previously unknown or sparsely studied heterogeneously expressed gene sets including metabolic pathways (amino-acid metabolism, carbon metabolism, siderophores) and physiological states (chemotaxis and motility). Interestingly, arginine biosynthesis genes appear to be heterogeneously expressed in both *E coli* and *B subtilis* in minimal media. Arginine synthesis is also isolated spatially (at the organ and cellular location level) in multicellular, eukaryotic organisms^28^. This conserved partitioning of a biochemical process suggests that some processes may be inherently more prone to segregate. We speculate that such segregation may be a result of incompatibilities between certain biochemical reactions and may have played a role in the initial development of multicellular structure.

Along with the method we describe in this work, two other groups have separately introduced a high-throughput bacterial scRNA-seq approach that is based on combinatorial indexing^29,30^. A commonality between these studies and ours is the discovery of subpopulations of cells expressing genes for specialized function, including genetic competence. While the different techniques uncover common biology, each has advantages and disadvantages. Our approach requires an upfront investment in probe generation and prior understanding of the genome, and is therefore not intended to be used with novel or environmental organisms that are not well characterized. Conversely, once a probe set is synthesized, our tool is fast, cheap, and provides very deep coverage of transcripts as well as a more accurate method of quantification by UMI. In addition, the technique relies on commercial equipment that is now ubiquitous in laboratories and core facilities around the world. Critically, unlike the combinatorial indexing system, our in-situ hybridization technique ensures that sequencing is not wasted on ribosomal or other non-mRNA transcripts, which account for between 93-97% of all transcripts that are sequenced by combinatorial indexing (ref).

Our technique of combining in-situ hybridization with microfluidic single cell partitioning also invites several potential uses for eukaryotic samples which often rely on targeting the 3’ poly-A tail; an approach that provides only a single site for transcript capture, excludes sequences lacking a poly-A, and provide limited information on splice variants. In addition, hybridization techniques are compatible with formalin-fixed paraffin-embedded tissues, the most common form of clinical sample, which are difficult to analyze using existing high-throughput single cell techniques. Finally, the extension of these techniques to mammalian and other eukaryotic cells could potentially be used to simultaneously identify the gene expression profile of an intracellular bacteria and its eukaryotic host cell.

In the context of microbial analyses these new tools will enable investigation of several recently observed phenomena in bacterial cultures. In particular, we foresee their use in characterizing bacterial subpopulations that are differentially resistant to antibiotics (antibiotic persistence), and the physiological role of bacterial cells that differentially express toxin and virulence genes in bacterial infections. In eukaryotic systems, where mature scRNA-seq tools have existed for several years, computational methods extend beyond surveys of the different cell types to provide information about the dynamics of cellular populations such as pseudotime arrangements of cells ^31^, lineage and causality maps and cell-type specific QTLs^32^. In bacterial communities we predict that these bioinformatic approaches will be used to understand dynamic processes like phage-bacteria interactions, the organization of biofilms, and lab-based evolution of traits.

## Acknowledgements

DS was funded in part by the Natural Sciences and Engineering Research Council of Canada (NSERC PGSD2-517131-2018). DS and SH acknowledge funding from NIH NIGMS R00GM118910, NIH NHLBI R01HL158269, U19 Systems Immunology Pilot Project Grant at Harvard University, and the Harvard University William F. Milton Fund. AZR and RAM were funded by a Dupont Science and Innovation award. Portions of this research were conducted on the O2 High Performance Compute Cluster, supported by the Research Computing Group, at Harvard Medical School. See http://rc.hms.harvard.edu for more information.

## Methods

### Strains and Growth Conditions

*Bacillus subtilis* str. 168 was used for all single cell Bacillus experiments. For experiments on suspension cultures in glucose/malate, cells were grown at 37°C with vigorous shaking in M9 media supplemented with CaCl2 (0.1 mM), 0.2% glucose, tryptophan, and trace metal mix ^8^. Trace metal solution was made as a 100X concentrate using Na-EDTA 5.2 g, FeSO_4_-7H_2_O 2100.0 mg, H_3_BO 330.0 mg, MnCl_2_-4H_2_O 100.0 mg, CoCl_2_-6H_2_O 190.0 mg, NiCl_2_-6H_2_O 24.0 mg, CuCl_2_-2H_2_O 2.0 mg, ZnSO_4_-7H_2_O 144.0 mg, Na_2_MoO_4_-2H_2_O 1200.0 mg, DI Water, 999.0 ml. The solution pH was adjusted to 7.0. At OD of 0.4-0.5 cultures were supplemented with 50 mM malate.

*Bacillus subtilis* strain PY79 was used for all fluorescent promoter-reporter strains. comG reporter strain was previously described ^8^. Promoter reporters for cotY and argC were made as previously described ^8^. Briefly, YFP promoter reporters were cloned into the ECE174 backbone plasmid which uses sacA integration site and encodes chloramphenicol resistance (R. Middleton, obtained from the Bacillus Genetic Stock Center). Strains were made by genomic integration into the genome. Fluorescent reporters were integrated into the sacA site and checked by sequencing. Reporter strain detail is found in Supplementary Table 4. Reporter strains were grown in the same media and growth conditions used in single cell experiments.

*E. coli* MG1655 was used in all *E coli* single cell experiments. For experiments in minimal media cells were grown overnight in M9 minimal media and incubated at 37°C with moderate shaking (200 rpm). Overnight cultures were diluted and subcultured and a fixed specimen was collected at mid exponential phase (OD 0.3-0.5). For experiments in anaerobic LB media cells were inoculated into degassed LB that was aliquoted inside a COY anaerobic chamber containing (95% N2 5% H2). Overnight culture was subcultured into a degassed media tube using a needle and syringe purged in N2 gas, and the samples were incubated anaerobically inside an anaerobic box with a gas-pack (manufacturer) at 37°C. Cells were fixed at OD 0.3-0.5 by addition of formaldehyde using a N2-purged syringe and needle directly into the tubes.

### Probe set design

Probes were designed with an mRNA complementary region approximately 50 bp in length flanked by a PCR handle (18-23 bp) towards the 5’ end and a 30 poly(dA) tract on the 3’ end. On the ends of each probes we included a region allowing for circularization and cutting with hindIII as outlined by Schmidt et al ^28^. Probe sequences are included in Supplementary Tables 1 and 2.

### Probe amplification

Probes were amplified using a rolling circle method similar to the one used by Schmidt et al ^28^ with slight modifications. To amplify our probes, we did not use nicking enzymes but instead only used the HindIII digestion site in the rolling circle scheme ^28^. Our incubations were scaled by a factor of 5 from the detailed protocol provided by Schmidt et al. Otherwise, all other aspects of the protocol were as previously detailed. Amplified probes were eluted and purified by PAGE electrophoresis using 15% urea gels and ethanol precipitation as described in Sambrook et al ^59^.

### UMI and Poly-A Addition

Amplified “proto-probes” were extended to include a unique molecular index (UMI) and 3’ poly-adenine tail by isothermal extension with a 3’ blocked primer containing the reverse complementary sequences. 100 ul of probes and 1 mg blocked extension primer (Supplementary Table 3) were mixed with 100 units of klenow fragment and 1x NEB buffer 2 and incubated for 30 min at room temperature. The extended oligonucleotide library was selectively purified by PAGE electrophoresis (final probe length approximately 142 bp) using 15% urea gels followed by ethanol precipitation as described in Sambrook et al. ^59^

### Fixation and in situ hybridization reactions

Cells from one intact colony or 2 mL of cell culture were fixed using a 30 minute incubation in 1% paraformaldehyde (final concentration) at room temperature. Formaldehyde-fixed samples were washed with 0.02 % SSC by gentle centrifugation (6,000 x G) for 2.5 minutes. After the wash, cell pellets were resuspended in 1 mL MAAM (4:1 V:V dilution of methanol to acetic acid). Samples were kept at −20°C for up to 2 days before further processing. For in-situ probe hybridization, 150ul of fixed sample was centrifuged and washed once in 1x PBS to remove methanol and acetic acid. After the wash step cells were incubated in 200 ul PBS with 350u/ul of lysozyme solution (Epicentre ready-lyse, Epicentre Madison WI USA) for 30 minutes at room temperature. After 30 minutes cells were pelleted and washed with 500 ul PBS-tween (1 x PBS with 0.1 % Tween 20). Cells were then re-suspended with 100 ul of probe binding buffer consisting of 5x SSC, 30% formamide, 9mM citric acid (pH 6.0). 0.1% Tween 20, 50 ug.ml heparin, and 10% low MW dextran sulfate) ^60^. Cell suspensions were placed in a 50°C shaker-incubator and allowed to pre-equilibrate for 1 hour. After an hour 50 ul of probes, (600 ng/ul) were added to each cell suspension, and samples were left to incubate overnight.

The next day samples were washed 5 times in pre-warmed (50°C) probe-wash solution (5 x SSC, 30% formamide, 9 mM citric acid (pH 6.0), 0.1% Tween 20, and 50 ug/mL heparin).

Before flowing on the 10X device cells were washed 3 times in 1x PBS and diluted as suggested in the 10X Chromium instruction manual.

### Microfluidic encapsulation and droplet generation reaction

Single cell partitioning, barcoding, and cDNA library generation was achieved using 10X Genomics’s Chromium Controller with the Chromium Single Cell 3’ Reagents Kit (v2 chemistry) as described by 10X Genomics (https://support.10xgenomics.com/singlecell-gene-expression/index/doc/user-guide-chromium-singlecell-3-reagent-kits-user-guide-v2-chemistry). The protocol was modified to achieve bacterial scRNA-seq. During GEM generation, a master mix containing the following reagents (per rxn, not accounting for excess volume) was prepared: 33 uL of 4X ddPCR Multiplex Supermix (BioRad), 4 uL of custom primer (10 uM), 2.4 uL additive A (10X Genomics), 26.8 uL dH2O. All other reagents specified by 10X Genomics were omitted.

Prepared cell samples were washed 3X in PBS and diluted to 1000 cells/uL before loading on the microfluidic chip (“Chip A Single Cell”) with a targeted cell recovery of 10,000 cells.

### Library construction & sequencing

After running the Chromium Controller, each sample (i.e. “reaction”) was visually inspected to confirm successful GEM formation (should observe a well distributed emulsion with a volume of approximately 100 uL). Samples were then transferred to fresh PCR tubes and cycled at the following conditions (replacing the “GEM-RT Incubation Step” in 10X Genomics protocol): 94°C for 5 minutes, 6 cycles of (94°C for 30s followed by 50°C for 30s then 65°C for 30s), hold at 4°C.

After PCR #1, the emulsion was broken and the pooled DNA purified using Dynabeads MyOne Silane as described in 10X Genomics’ protocol. Purified DNA was amplified once more (replacing the “cDNA Amplification step” of 10X Genomics’ protocol) using a master mix composed of 17 uL of purified DNA, 20 uL of Q5 Hot Start 2X MM, 1.5 uL of forward primer (10 uM) and 1.5 uL of reverse primer (10 uM). PCR conditions included a 30s incubation at 98C followed by 16 cycles of 10s at 98C, 20s at 62C, and 20s at 72C before a final extension at 72C for 2 min.After PCR #2, amplified DNA was purified using the NucleoSpin® Gel and PCR Clean-up kit (Macherey-Nagel) as per manufacturer’s instructions. Purified DNA was run on Agilent TapeStation to confirm the presence of a band at 180 bp, indicating successful library generation. Libraries then prepared for sequencing by Illumina adapter addition via low cycle (n = 6) PCR with custom library preparation finishing primers (see Supplementary Table 3).

Sequencing was done on an illumina nextSeq-1000 instrument with 100 cycle reagents. Libraries were spiked with 30% phi-X and sequenced for 8bp in the i7 index direction and 119 bp for “Read 1”. Addition of 30% phiX improved fastQ quality scores.

### Microscopy

Cultures were visualized on a Leica DM3000 light microscope before single cell experiments to ensure samples were free of chains and clumps. Reporter strains were imaged using a 100X oil-immersion objective on a Zeiss Axio Observer inverted microscope equipped with a colibri-7 LED fluorescent light source and an axiocam digital camera. Flagellar staining was done using the Remel flagellar stain (Thermo-Fisher Scientific) as per manufacturer instructions.

### Image analysis using cell segmentation and outlier identification

Cells in microscopy images were segmented using the microbeJ program within FIJI (ImageJ). Outliers were identified by using the IQR outlier method. Specifically, quartiles and the interquartile range (IQR) were determined for each dataset. Outliers were determined by standard outlier detection parameters: cells that were more than 1.5 IQRs above Quartile 3 were designated as cells within an overexpressing population. Population histograms were plotted and outliers reported.

### Constructing a Reference Genome

Here we describe how we constructed reference genomes tailored to the probesets used for each organism (see Methods Section <u>Probeset Design</u>). For a given organism, let *P* be the number of probes in the probeset 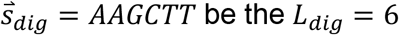 be the *L*_*dig*_ = 6 base pair (bp) nucleotide sequence of the HindIII digestion site (see Supplementary Figure S1a), 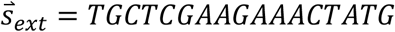 be the *L*_*ext*_= 17 bp nucleotide sequence of the extender, 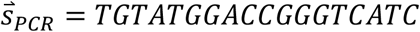 be the *L*_*pcr*_ = 18 bp nucleotide sequence of the PCR handle, and 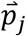 be the *L*_*j*_ bp nucleotide sequence of the *j*^th^ probe. Let 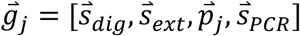 be the *L*_*dig*_ *+ L*_*ext*_*+ L*_*j*_ *+ L*_*pcr*_ bp nucleotide sequence formed by flanking the *j*^th^ probe by the HindIII digestion site, extender sequence, and PCR handle as shown in Figure 1a. We first constructed a FASTA file by concatenating all *p* of the 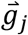 sequences. Next, we constructed a GTF file to index the FASTA. Typically, the GTF indexes genes in the FASTA by recording the location of the starting and ending bps of each gene. Since we have multiple probes per gene, and our FASTA is a concatenation of these probes, we therefore use the GTF to index probes instead. Specifically, we recorded that for the *j*^th^ probe, the starting and ending bps of 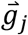 in the FASTA are 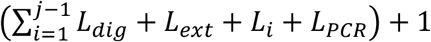 and 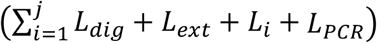 respectively. We use 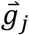 as the definition of a probe instead of 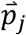 for the purpose of alignment because sequenced reads should contain the flanking sequences anyways so including them minimizes spurious alignment of off-target reads. Lastly, we constructed a reference genome by using the *mkref* command in CellRanger (v3.1.0), supplying the generated FASTA and GTF files to the *--fasta* and *--genes* arguments respectively.

### Creating FASTQ Files for CellRanger

Each read in the Illumina-sequenced FASTQ file contains the 10x CB, the 10x UMI, the Probe UMI as well as the mRNA sequence on a single line. This file therefore needs to be re-formatted to be compatible with CellRanger, so that the CB and UMI is stored in the R1 file and the mRNA sequence is stored in the R2 file. We also perform filtering at this stage so that only reads containing all the information needed by the CellRanger pipeline are retained. First, we discard reads whose length is less than or equal to 110 bp. Next, we aligned each read to 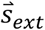 using the *nwalign(glocal=true)* function in MATLAB (2019a), and discarded reads where the alignment score was <17. We search for the extender sequence specifically, because for all probe designs the Extender sequence flanks the Probe UMI sequence (see Supplementary Figure S1c).

We create two sets of FASTQ files using reads retained after these filtering steps. In the first set of FASTQ files, we record the 10x CB and Probe UMI in R1, with the Probe UMI obtained by extracting 12 bps in either the 5’ or 3’ direction from the Extender sequence according to the probe design (see Supplementary Figure S1c). In the second set of FASTQ files, we record the 10x CB and 10x UMI in R1. For both sets of FASTQs, we trimmed all nucleotides 5’ (3’) of the Extender sequence if the Extender is 5’ (3’) of the binding region, and record the result in R2. The two sets of FASTQs therefore have the same number of lines, identical R2 files and the same 10x CBs in the R1 files, and only differ in the UMIs recorded in the R1 files. We obtained single cell count matrices from these reformatted FASTQs by using the *count* function in CellRanger (v3.1.0) and supplying the custom made genome (described in Methods Section <u>Constructing a Reference Genome</u>) as the *--transcriptome* argument. Since CellRanger discards reads if QC scores for UMIs are too low, the same set of reads may not necessarily end up being used in constituting single-cell count matrices from the two sets of FASTQ files.

### Comparing Single cell Probe Expression Matrices Generated Using 10x vs Probe UMIs

The output from CellRanger is a single cell matrix where each row, *i*, corresponds to one of *N* cells, and each column *j* corresponds to one of *p* probes. When the supplied FASTQ files have Probe (10x) UMIs recorded in R1, each entry in the single cell matrix, 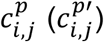 is the number of Probe (10x) UMIs in cell *i* corresponding to probe *j*. The Probe UMI remains fixed over multiple rounds of PCR and can therefore be treated as a classical UMI, while in our protocol, new 10x UMIs may be introduced on the same transcript at each PCR round. Therefore, 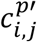 may not accurately record the number of transcripts. We used 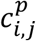 for all results presented in the main text of the paper. Since it may be more convenient to use 10x UMIs directly, we also quantified the extent to which 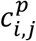 differs from 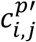 by computing a joint distribution of count frequencies, 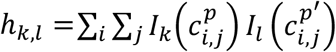, where *I*_*s*_(*t*) = δ_*s,t*_ (see histograms in Supplementary Figures S6-S8).

On the one hand, when *h*_*k,l*_ > 0, for *l* > *k*, there is over-counting; the sequencing depth is large enough to detect spurious 10x UMIs introduced in latter PCR rounds. On the other hand, when *h*_*k,l*_ > 0, for *l* < *k*, there is under-counting; CellRanger detects that the same 10x UMI is coincidentally found on different probes, and discards all such reads. Since under-counting doesn’t introduce spurious transcript counts and can be treated in the same vein as low sequencing depth or low capture efficiency, we do not consider it a confounder. The accuracy of a protocol relying only on 10x UMIs can therefore be quantified by the fraction of 10x UMIs corresponding to entries in *h*_*k,l*_ with *l* ≤ *k*:

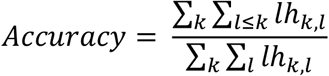

For the *B. Subtilis* sample, the accuracy of using 10x UMIs thus computed was only 41%. However, when we computed gene expression matrices using 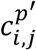 (see Methods Section <u>Single cell Gene Expression Matrices from Single cell Probe Expression Matrices below</u>) and performed differential gene expression (see Methods Section <u>Transcriptomic Analysis and Visualization</u> below), we were still able to detect clusters corresponding to competence and sporulation (see e.g. clusters #2 and #11 respectively in Supplementary Figure S6 and Supplementary Table 7). Likewise, for the *E. coli* sample grown in minimal media, the accuracy was only 33%, but we were still able to detect clusters corresponding to the *fim* operon, cell motility and differential usage of carbamoyl phosphate (see e.g. clusters #8, #2 and #7 respectively in Supplementary Figure S7 and Supplementary Table 7). Lastly, for the *E. coli* sample grown in LB media, the accuracy was 88%. Here too, we were able to recapitulate biological results obtained using Probe UMIs including clusters with different metabolic programming and upregulation of the tdc operon (see clusters #6 and #9 respectively in Supplementary Figure S8 and Supplementary Table 7). Taken together, this suggests that users of our technology may be able to use 10x UMIs directly i.e. 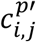. However, for the remainder of this study we restrict our attention to 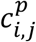.

### Single cell Gene Expression Matrices from Single cell Probe Expression Matrices

In order to perform standard single cell analysis such as cell-similarity clustering and differential gene expression, we need to determine 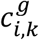, the number of transcripts of gene *k* in cell *ii*, whereas the output from CellRanger is 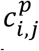. Let *G*_*k*_ be the indices of probes in the probeset corresponding to gene *k*. Since multiple probes may bind to the same transcript, setting 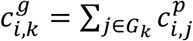 may result in an over-estimate of transcript counts. To be conservative while also retaining as many UMIs as possible, we therefore picked 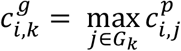 which lower bounds gene expression. This choice resulted in 66%, 79% and 81% of the UMIs in *c*^*p*^ being retained in *c*^*G*^, accounting for 65%, 77% and 81% of the total reads, for the *B. Subtilis, E. Coli* in minimal media and *E. Coli* in LB media samples respectively.

That different probes may be selected to represent the same gene in different cells is not material, since probes are selected proportional to their binding affinity. However, this choice introduces a potential bias since the gene expression tabulated in *c*^*G*^ will tend to be higher for genes with more corresponding probes in the probeset, i.e. larger |*G*_*k*_ |. As a sanity check, we also tabulated single cell gene expression matrices using 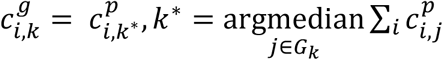 i.e. using a fixed probe across all cells for each gene corresponding to the probe with the median bulk expression. This choice only resulted in 10%, 18% and 17% of the UMIs in *cc*^*pp*^ being retained in *c*^*G*^, for the *B. Subtilis, E. Coli* in minimal media and *E. Coli* in LB media samples respectively. Despite the decreased transcriptional resolution, we are able to still detect clusters corresponding to the results presented in the main text (see Supplementary Figures S9-S11 and Supplementary Table 8). We therefore defer investigation of more sophisticated methods of combining probe counts to form a single cell gene expression matrix to subsequent work.

### Transcriptomic Analysis and Visualization

Here we describe how we analyzed single cell gene expression matrices, *c*^*g*^, using Seurat (v3.1.2 with default parameters except where indicated). We first log-transformed the data using the *NormalizeData(normalization*.*method = “LogNormalize”, scale*.*factor = 10000)* function, and selected the 2000 most variable genes using *FindVariableFeatures(selection*.*method = “vst”, nfeatures = 2000)*. Then, we z-scored these highly-variable genes using *ScaleData*. Next, we performed linear dimensionality reduction using PCA down to 50 dimensions (*RunPCA*). Points in this embedding were used to construct UMAP plots (*RunUMAP(dims=1:10)*) and find neighbors for clustering (*FindNeighbors*). *FindClusters(resolution=1*.*0)* was used to run the Louvain clustering algorithm and generate clusters. We confirmed that the clusters highlighted in the main text appeared consistently for a range of resolutions from 0.5 to 1.5. Summary statistics of the number of genes/transcripts in a cell and cluster populations for the different samples can be found in Supplementary Table 5. Differential gene expression was performed using the *FindMarkers(min*.*pct=0*.*25)* function with a log-fold change cutoff of 0.25, whose output when applied to our data is shown in Supplementary Tables 6-8. Heatmaps of differentially expressed genes were generated using the *DoHeatMap* function, and show the top over-expressed genes in each cluster with Bonferroni-corrected p-value < 0.05.

## Data and Code Availability

All data used in this paper will be made available on the NCBI GEO upon publication. Code to prepare custom references, reformat FASTQ files for the CellRanger pipeline and generate single-cell gene expression matrices from single-cell probe expression matrices is available at https://gitlab.com/hormozlab/bacteria_scrnaseq. All code and instructions to reproduce numerics and figures in the manuscript will be made available upon publication.

## Supplementary Figures

**Figure S1:** Detailed schematic of C2C-based proto-probe amplification and primer extension incorporation of probe-based UMI and polyA

**Figure S2:** Heatmap of *B subtilis* cells from Figure 3 including all marker gene names

**Figure S3:** scRNA-seq of *B subtilis* using in-droplet reverse transcription (RT) instead of in-droplet PCR identifies sporulation and competence populations

**Figure S4:** Heatmap of aerobic M9 culture of E coli cells from Figure 4 including all marker gene names

**Figure S5:** Heatmap of *E coli* fermenting in LB media from Figure 4 including all marker gene names

**Figure S6:** scRNA-seq of *B subtilis* using 10x UMIs instead of Probe UMIs

**Figure S7:** scRNA-seq of *E coli* cells in aerobic M9 culture using 10x UMIs instead of Probe UMIs

**Figure S8:** scRNA-seq of *E coli* cells in LB media using 10x UMIs instead of Probe UMIs

**Figure S9:** scRNA-seq of *B subtilis* using bulk median instead of per-cell maximum probe counts for gene expression

**Figure S10:** scRNA-seq of *E coli* cells in aerobic M9 culture using bulk median instead of per-cell maximum probe counts for gene expression

**Figure S11:** scRNA-seq of *E coli* cells in LB media using bulk median instead of per-cell maximum probe counts for gene expression

**Figure S12:** Frequency of marker genes and spores in the population measured by microscopy

## Supplementary Tables

**Table S1:** *B subtilis* probes ordered from TWIST Bioscience

**Table S2:** *E coli* probes ordered from TWIST Bioscience

**Table S3:** Probe amplification reagents and costs

**Table S4:** Strains used in this study

**Table S5:** Sample summary

**Table S6:** DGE analysis

**Table S7:** DGE analysis using 10x UMIs instead of Probe UMIs

**Table S8:** DGE analysis using bulk median instead of per-cell maximum probe counts

**Table S9:** Genes in each section of the Venn diagram in Figure 3d

**Table S10:** Genes used in volcano plots on Figure 3C and Figures S2, S4 and S5

## Supplementary Notes

**Supplemental note 1:** Additional biology related to Figure 3 and Figure S2

**Supplemental note 2:** Additional info on Figure S3 and use of droplet RT instead of PCR

**Supplemental note 3:** Additional biology related to Figure 4a and Figure S4

**Supplemental note 4:** Additional biology related to Figure 4b and Figure S5

**Supplemental note 5:** Calculation of probe concentrations

---

**Supplemental note 1:** Additional biology related to Figure 3 and Figure S2

For *B. subtilis* in minimal media, in addition to the competent and sporulating cell subpopulations discussed in the main text, we observe two other clearly distinguishable cell groups indicative of distinct biological states. The largest group of cells (comprising clusters 1, 2, 3, 5, and part of cluster 7) upregulate dhbA, dhbB, dhbC, dhbE, and dhbF (the 5 enzymes implicated in the biosynthesis of bacillibactin) as well as yusV, an ABC transporter of bacillibactin. In addition, at least 3 out of the 5 clusters within the subpopulation showed upregulation of genes involved in thiamine biosynthesis (tenA, tenI, thiD, thiG, thiF, thiS, thiO; gene set enrichment = 16.85, FDR = 2.79E-05), amino acid biosynthesis (fold enrichment = 5.23, FDR = 1.48E-07), amino acid activation (thrS, gatB, ileS, aspS; fold enrichment = 6.22, FDR = 3.46E-02), surfactin biosynthesis (srfAA, srfAB, srfAC, srfAD), “de novo’’ IMP biosynthesis (gene set fold enrichment = 25.79, FDR = 9.10E-09), purine biosynthesis (purB, purC, purD, purQ, purH, purF, purL, purK, purM, purS; gene set fold enrichment = 21.66, FDR = 1.83E-02), pyrimidine biosynthesis (pyrAA, pyrK, pyrF, pyrH; gene set enrichment = 9.63, FDR = 2.59E-02), chorismate biosynthesis (aroA, aroB, aroC, aroH, aroE; gene set fold enrichment = 18.05, FDR = 4.60E-03), cell motility (hag, fliY, fliD, fliH, flhP; gene set enrichment = 6.19, FDR = 1.44E-02), sulfate assimilation (cysI, cysH, cysC, cysJ; gene set enrichment = 18.05; FDR = 4.56E-03), and biotin biosynthesis (bioA, bioB, bioD, bioF, bioW; gene set fold enrichment = 18.05, FDR = 4.65E-03). Altogether, this subpopulation appears to represent the most metabolically active cell state within the total population, an observation further supported by the fact that genes encoding ribosomal components & translation machinery were also enriched (rpmI, rplU, rplR, rpsM, rpsE, rpsI, rpsN, rpsD, rpoB, rplV, rplX, rplB, rpmA, rpsS, rplS, rpsP, rplL, rpmD, rpmC; “translation” gene set fold enrichment = 7.81, FDR = 3.07E-12). Cell cluster #2, part of the subpopulation, may represent the gateway into sporulation as early sporulation factors sigF, spooA, and spoIIAB are all uniquely upregulated and cells are projected in close proximity with cluster 9 after dimensional reduction. In addition, these cells downregulate genes implicated in motility (hag, fliD, fliG, fliY, fliK, fliM, flgK, flgL) in stark contrast to other cells within the subpopulation. Furthermore, cells within cluster #5 are distinguished from others in the “metabolically active” subpopulation by expression of genes related to arginine biosynthesis via ornithine. L-arginine is used by cells for protein synthesis as well as for production of polyamines such as putrescine and spermidine; compounds which have functions in nucleic acid binding, protein activation, and membrane stabilization due to their polycationic state. The 4^th^ distinct cell state detected in *B. subtilis*, only slightly less in size compared to the “metabolically active” cell group (45% of total population), was characterized by lower expression of many of the genes upregulated in the “metabolically active” subpopulation. Uniquely, cells in cluster 0 (930 cells, 76% of the subpopulation) were found to overexpress sdpB (logFC = 0.68, adjusted p-value = 4.7E-4). Under starvation conditions, *B. subtilis* produces an antimicrobial peptide through the sdpABC operon that is active against its own species, a tactic used by the cell to increase its own chance of survival and delay commitment to sporulation. While the sdpC gene encodes the 42AA toxic peptide, sdpA & sdpB are required for maturation of the peptide into its active form prior to secretion. sdpB is a multipass membrane protein which affects the toxicity of the final SDP peptide without being involved in signal peptide cleavage, disulfide bond formation, or secretion. The sdpABC operon was found expressed in cells within each of the four subpopulations, however, it is only within cluster #0 that sdpB was found to be significantly upregulated. This finding, along with the overall reduction in transcript load as compared to the “metabolically active” subpopulation, suggests a more dormant biological state in which cells appear to be facing a nutritional strain but have not yet committed to an alternative developmental program. This hypothesis is supported by the fact that cells in cluster 7 are split between the two largest subpopulations in the UMAP projection as well as by the clustering of a few cells associated with competence (part of cluster 8) near this dormant subpopulation – possibly suggesting a developmental transition between the two states.

**Supplemental note 2:** Additional info on Figure S3 and use of droplet RT instead of PCR

While the approach in Fig S3 (RT in drops instead of PCR) proved less sensitive in transcript capture compared to the final, optimized method, we still resolve distinct biological states upon dimensional reduction of expression vectors including cell subpopulations corresponding to competence and sporulation. Additionally, we observe a majority grouping of cells (clusters #0-5) with significantly increased expression of ribosomal genes as well as genes involved in purine and amino acid biosynthesis – corresponding to the “metabolically active” subpopulation defined previously. scRNA-seq also revealed a subpopulation of cells (cluster #8, 1.9% of cells) within the sample exhibiting a state unforeseen in the previous run - characterized by the upregulation of genes encoding a manganese importer (mntABCD, logFCs > 1.99, adjusted p-values < 1E-11), sigma factor sigW (logFC = 1.69, adjusted p-value = 5E-16), and stress response proteins (yuaF, yceDEF, yqfA, yqeZ, logFCs > 0.99, adjusted p-values < 3.3E-04) involved in maintaining cell membrane integrity. yceF, specifically, is a cell envelope stress protein that conveys resistance to manganese. katA, an iron-binding catalase which protects cells against toxification from hydrogen peroxide, was also upregulated. Altogether, this cluster of cells appears to be responding to stress induced by a low concentration of manganese within the environment by upregulating manganese uptake machinery. maeA was also upregulated in these cells (logFC = 1.9, adjusted p-value = 3.65E-34). maeE encodes a malic enzyme necessary for catabolic repression of pyruvate import (pftAB) as well as for maintaining ATP levels during growth on the carbon source by conversion of malate to pyruvate and generation of NADH.

**Supplemental note 3:** Additional biology related to Figure 4a and Figure S4

Single cell transcriptomes of *E. coli* grown in M9 were clustered into ten groups. In the largest group of cells (cluster #0, 514 cells, 15.5% of total), ribosomal genes (rpl operon, rpsC, rpsD, rpmC, rpmD) as well as deaD and priB are amongst the most upregulated compared to the rest of the cell population. deaD (logFC = 0.7, adjusted p-value = 2.3E-11) is a DEAD-box RNA helicase involved in ribosome biogenesis, translation initiation, and mRNA degradation at low temperatures - forming a cold-shock degradosome with RNase E. Kuchina et al. also identified a subpopulation of *E. coli* differentially expressing deaD, along with other cold shock proteins, in their scRNA-seq experiments and hypothesized the cluster was an artifact of sample preparation. Since our protocol fixes cells directly in culture, we find it unlikely that this state is a product of cold shock - a conclusion reinforced by the fact that we observe no other cold shock associated proteins within the set of upregulated genes. priB (logFC = 0.37, adjusted p-value = 6.4E-12) encodes a protein within the primosome which catalyzes lag strand priming during DNA replication. Conversely, genes involved in pyrimidine biosynthesis (pyrI, pyrB, pyrC) were significantly downregulated within the cluster. Interspersed with cluster #0 on the dimensionally reduced plot, we observe a group of cells (cluster #8, 141 cells) highly upregulating genes from the fim operon, as discussed in the main text. Cells in cluster #2 appear less active overall, as all genes found to be significantly differentially expressed within the cluster are downregulated compared to the rest of the population. This may be an effect of poor probe penetration within the protocol or indicative of a more basal cell state. Cluster #3 reveals no significantly overrepresented gene sets by GSEA after accounting for the false discovery rate. Upregulated genes reported in the cluster include ompT and ompX as well as tolB and tolC (logFCs > 0.25, adjusted p-values < 1E-14). In addition, scRNA-seq reveals a cluster of cells (cluster #4, 403 cells, 12% of total) that appear to be entering stationary phase; exhibiting a relative increase in expression of dps (logFC = 0.91, adjusted p-value = 4.17E-26). dps encodes a protective protein which binds to the bacterial chromosome and sequesters intracellular Fe2+, forming a highly stable protein-DNA mineral complex which protects the DNA from oxidative damage. In E. coli, dps has been associated with the transition into stationary phase as cells quickly work to preserve genetic material from stressors including UV irradiation, metal toxicity, thermal fluctuations, and acid/base shock. Cluster #9 (117 cells, 3.5% of total) was also shown to upregulate dps (logFC = 1.05, adjusted p-value = 7.04E-16) while highly expressing the gad (gadA, gadB, gadC) and hde (hdeA, hdeB, hdeD) operons. Both operons encode resistance to acid stress and fall under the gene set of “regulation of intracellular pH” (fold enrichment > 100, FDR = 6.72E-03). gadA and gadB produce the subunits of glutamate carboxylase which allows cells to interconvert between L-glutamate and GABA by incorporation of an intracellular proton while gadC encodes the antiporter enabling L-glutamate uptake and GABA export. hdeA and hdeB encode acid stress chaperone proteins with differing pH optimums; both preventing the aggregation of periplasmic proteins denatured under acidic conditions. osmY, encoding a periplasmic chaperone protein, was also upregulated within this cluster and has been shown to be induced by entry into the stationary phase. Altogether, DGE analysis reveals a fraction of cells within the population that are experiencing significant stress. Cells in cluster #4 appear to be at the initiation of this state while cells in the cluster #9 are more clearly induced, expressing dps along with proteins dealing with acid, osmotic, and oxidative stresses. It is possible these clusters represent cells transitioning into stationary phase or, alternatively, a transient state experienced during active metabolism/exponential growth. Cells within clusters 5 and 7 upregulate genes involved in pyrimidine biosynthesis as discussed in the main text. Interestingly, cluster #5 differentially overexpresses argG (logFC = 0.89, adjusted p-value = 6.57E-29), although no other genes specific to arginine biosynthesis were found to be upregulated. Both clusters upregulate ygjF, encoding a G/U mismatch-specific DNA glycosylase to correct mispairings in double stranded DNA that arise from alkylation or deamination of cytosine. ygjF is often active in stationary-phase cells 36. Cluster 6 is defined by the upregulation of genes involved in arginine biosynthesis as discussed in the main text.

**Supplemental note 4:** Additional biology related to Figure 4b and Figure S5

Upregulated genes within the largest cell cluster (cluster #0, 677 cells, 18% of total) included fliC (LogFC = 1.02, adjusted p-value = 3.81E-54), uncharacterized genes ygeWXY (logFC > 0.7, adjusted p-values < 3.3E-32), genes putatively involved in the conversion of adenine to guanine during nucleotide salvage (xdhA, xdhD, logFCs > 0.85, adjusted p-values < 1E-34), ssnA (logFC = 0.80, adjusted p-value = 9.8E-23), a putative, phase-dependent regulator of cell viability during later stage growth, and genes in the tna operon (tnaABC, logFCs > 0.45, adjusted p-values < 1E-28). tnaA encodes tryptophanase, which converts L-tryptophan to indole and pyruvate as part of amino acid catabolism while tnaB is a low affinity permease and tnaC is a positively regulating leader peptide. Cell clusters #1 and #4, adjacent and densely packed groups on the UMAP, appear to represent a subpopulation of cells responding to over-acidification of the cytoplasm. Both clusters significantly upregulate the gad operon (gadABC) as well as dps. In addition, acid stress chaperone proteins hdeA and hdeB are overexpressed in cluster #4. No statistically significant gene sets were identified from upregulated genes in clusters #2-3, however, cells in both clusters express genes related to anaerobic fermentation as expected under the culture conditions. Cells in cluster #5 (283 cells, 7.5% of total) are characterized by the upregulation of genes involved in aerobic respiration as discussed in the main text. We also observe the strong upregulation of genes involved in ribosome assembly (fold enrichment = 15.26, FDR = 6.16E-21) and translation machinery (fold enrichment = 12.01, FDR = 6.26E-33) in these cells, indicating a more active metabolic state which can most likely be attributed to the increases in ATP yield that is achieved by aerobic respiration. We observe similar enrichment of ribosome-related genes in cell cluster #6 (191 cells, 5% of total) along with the upregulation of a few genes in glycolysis (eno, lpd, aceEF, logFC > 0.5, adjusted p-values < 2.2E-06) and oxidative phosphorylation (atpD, cydAB, logFC > 0.7, adjusted p-values < 1E-06). These findings suggest that cluster #6 is in a differentiating state, transitioning between aerobic and anaerobic metabolism. Interestingly, cell cluster #7 (135 cells, 3.6% of total) appears to capture cells undergoing anaerobic metabolism while responding to oxidative stress. Gene sets describing anaerobic respiration (acnA, yjjI, nrfA, frdC, frdA, ynfF, ynfE, nrfB, dmsA; fold enrichment = 8.00, FDR = 5.63E-03) and oxidative stress (acnA, ahpF, osmC, qor, pgi, sufD, ydbK, yliT, yggE, clpA; fold enrichment =, FDR = 2.68E-02) were both identified as overrepresented in the list of upregulated genes within the cluster. In addition, stress proteins such as hspQ (a heat shock response protein) and osmY (hyperosmotic response protein) were upregulated along with flu (logFC = 0.47, adjusted p-value = 2.88E-14), a surface-display, self-recognizing adhesin protein encoding gene which enables auto aggregation and flocculation of cells in static liquid culture as well as biofilm formation by uropathogenic *E coli* in situ. Cluster #8 (111 cells, 3% of total) is characterized by the strong upregulation of the tdc operon (threonine metabolism under anaerobic conditions) as discussed in the main text.

**Supplemental note 5:** Calculation of probe concentrations

<u>Determining Probe Concentration</u>

To determine the concentration of probes needed to saturate an approximate number of transcripts in solution, saturation was defined as total, nonspecific probe coverage for each transcript. In other words, a pool of transcripts is considered saturated when, for each molecule, there is an entire library of probes (n = 1 probe per unique design) exclusively dedicated. In reality, this system would be over-saturated as it contains enough probes to tag every transcript even if all transcripts have the same sequence.

In our protocol, we prepare 150 uL of fixed cells from a 2X concentrated culture collected at mid-log phase. Assuming the culture contains 1E8 cells/mL prior to concentration and each bacterial cell contains 2,000 transcripts ^13^, the following estimate can be made using a probe set containing 20,000 unique ssDNA probes of length 138 bp (∼42,500 g/mol).

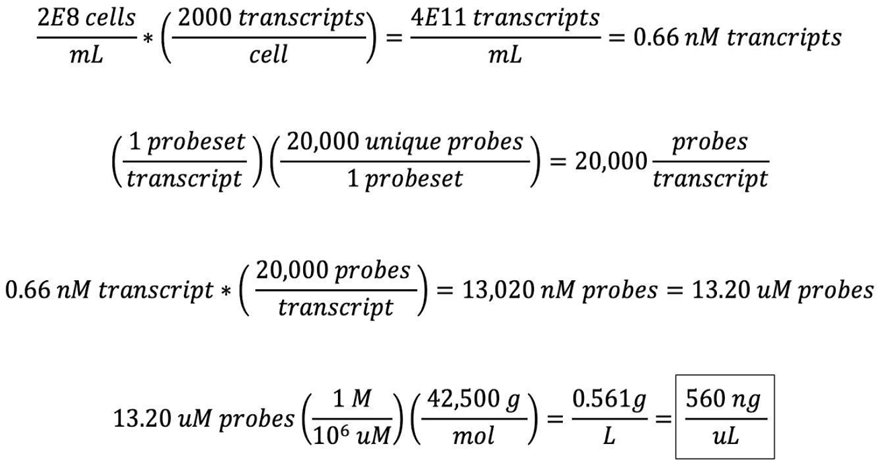

In our final experiments, we achieved a probe concentration of 600 ng/uL (0.6 mg/mL). To note, our calculations assume equal representation of all probe designs in the applied probe library which is unlikely due to biases introduced during probe synthesis and amplification. However, in all likelihood, this number represents a gross overestimate of probe requirements for a few reasons. Cell concentration and, therefore, transcript load was purposefully overestimated by assuming an exponentially growing culture contains 1E8 cells/mL; this number better represents the carrying capacity of *B. subtilis* in minimal media. In addition, our definition of “saturation” was overly strict, as discussed above. Lastly, probe libraries were designed redundantly to minimize noise in the downstream analysis by creating multiple probes to target different regions of the same gene. For the *B. subtilis* library, we designed, on average, 7 probes per gene - further assuring that our final probe concentration is sufficient to tag all available transcripts for scRNA-seq.

## Notes

https://gitlab.com/hormozlab/bacteria_scrnaseq

### Competing Interest Statement

The authors have declared no competing interest.

### Summary of Updates

Includes supplementary figures.

