## Supplementary Figures for "Droplet-based single cell RNA sequencing of bacteria identifies known and previously unseen cellular states"

**Figure S1:** Detailed schematic of C2C-based proto-probe amplification and primer extension incorporation of probe-based UMI and polyA

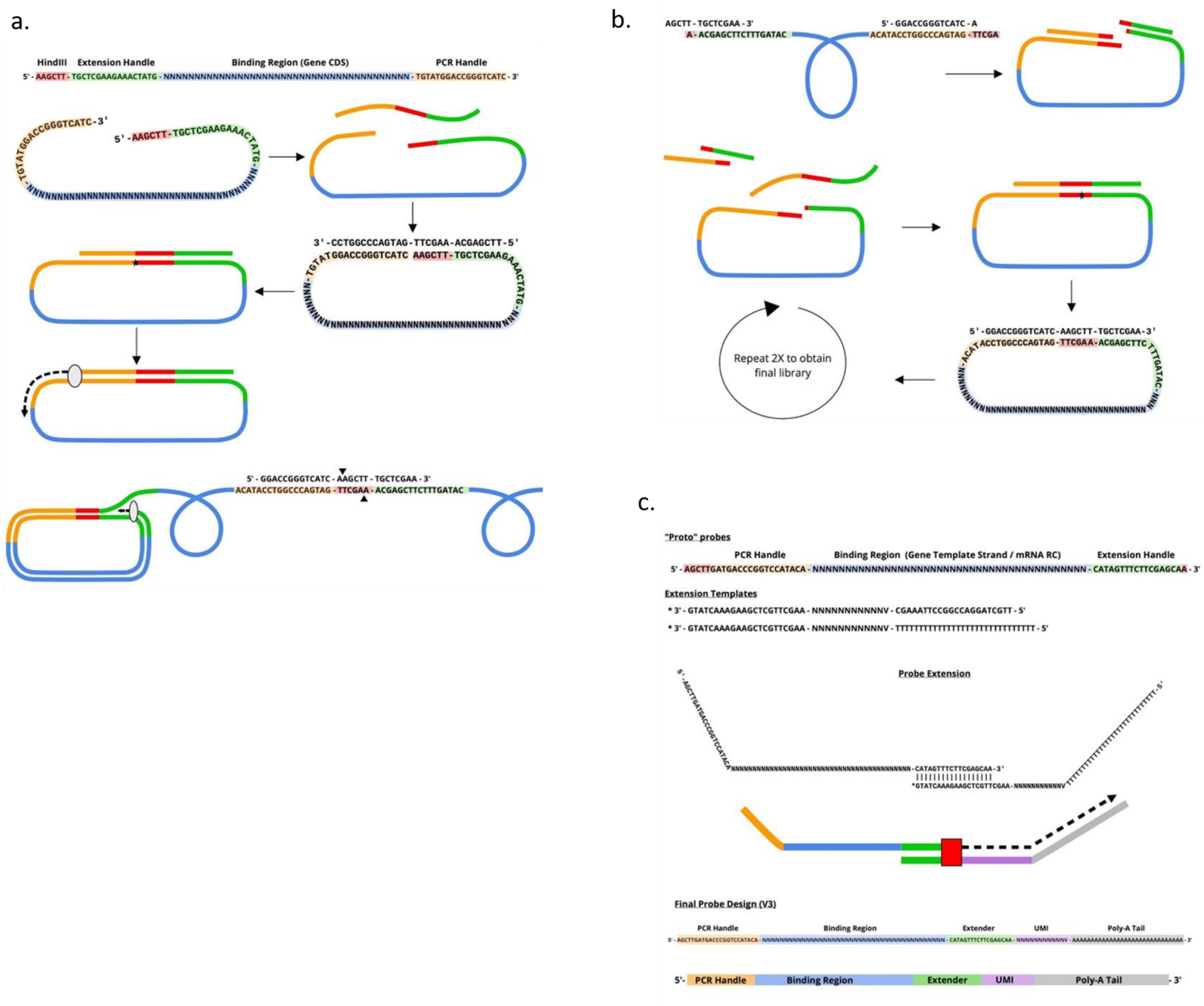


**a.** Oligonucleotide pools ordered from TWIST biotech were circularized by addition of a scaffolding primer that contained a sequence with reverse-complementarity to the two adjoining ends. Circles were finalized with T4 ligase and the scaffolding primer was used as a primer for phi29-mediated rolling circle amplification. **b.** Amplified circles were cut into single-strand primers by addition of nicking primer with reverse complementarity to the hindIII containing region and the two end-joints. Circles were made with these newly liberated single-strand amplified probes by adding the appropriate reverse-complement primer and ligation and the phi29 reaction was carried out as before. Libraries were amplified in 3 total rounds of rolling circle reactions. **c.** After digestion of the products from the third and final rolling circle amplification proto-probes were purified by gel electrophoresis and a UMI and polyA tail were added by primer extension using a blocked extension template.

**Figure S2:** Heatmap of B subtilis cells from figure 3 including all marker gene names

a.

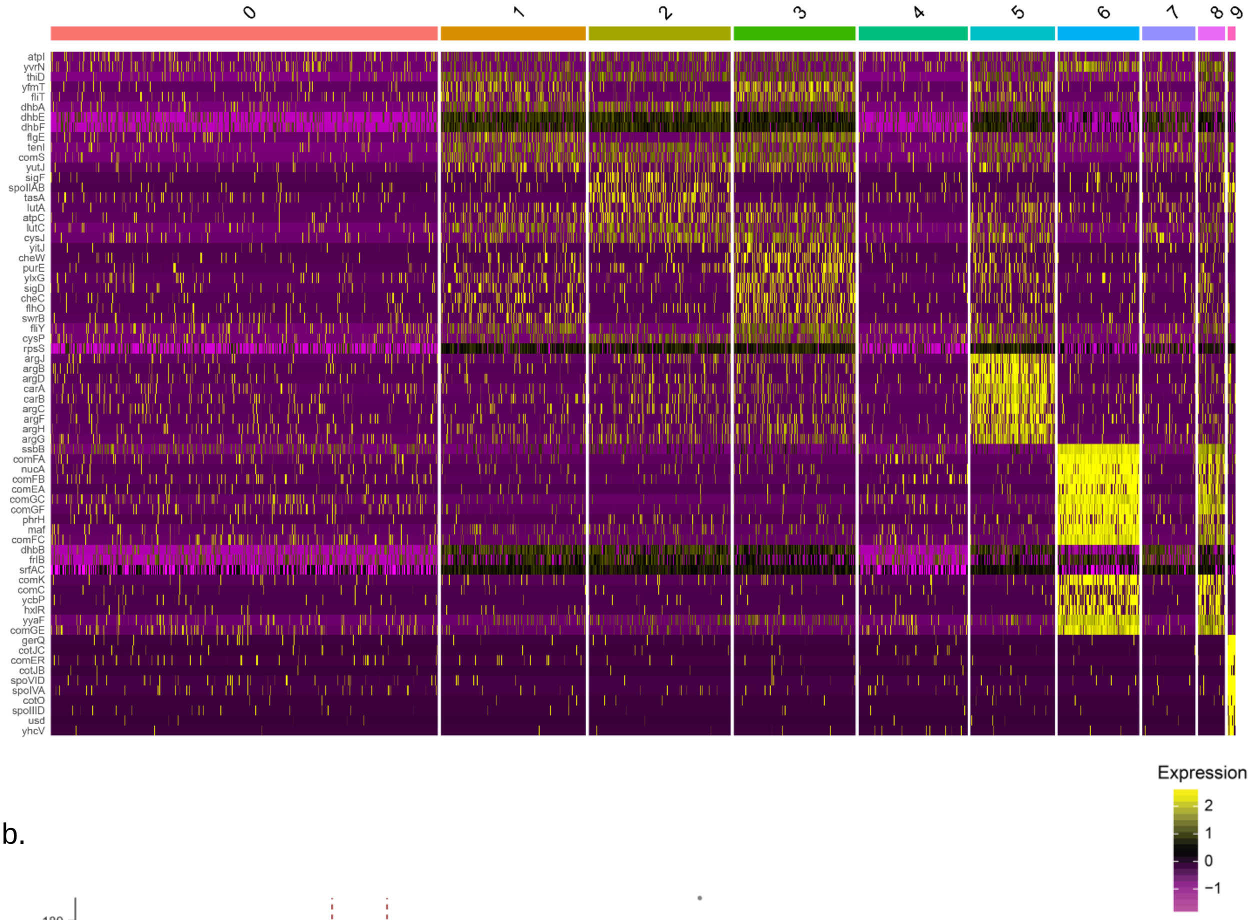


b.

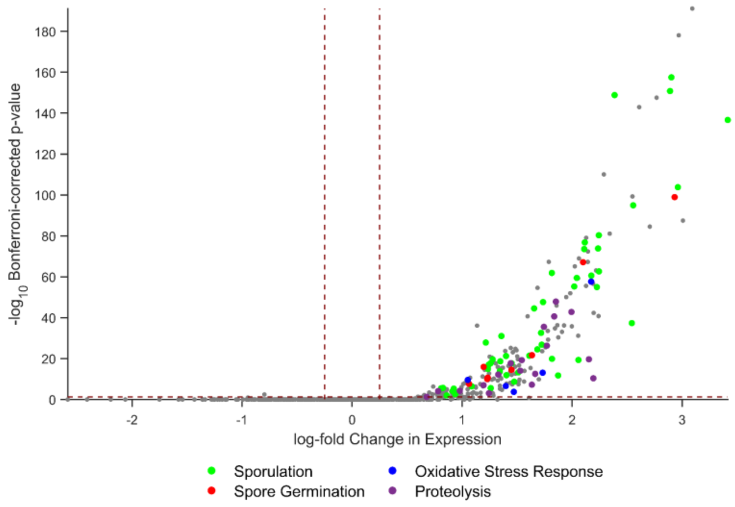


**a.** Heatmap as in figure 3a. including names for all marker genes
**b.** Gene enrichments in cluster 9 represent sporulation genes as in Figure 3c as well as other gene-sets enriched in this cluster and highlighted in this panel.

**Figure S3:** scRNAseq of *B subtilis* using in-droplet reverse transcription (RT) instead of in-droplet PCR identifies sporulation and competence populations

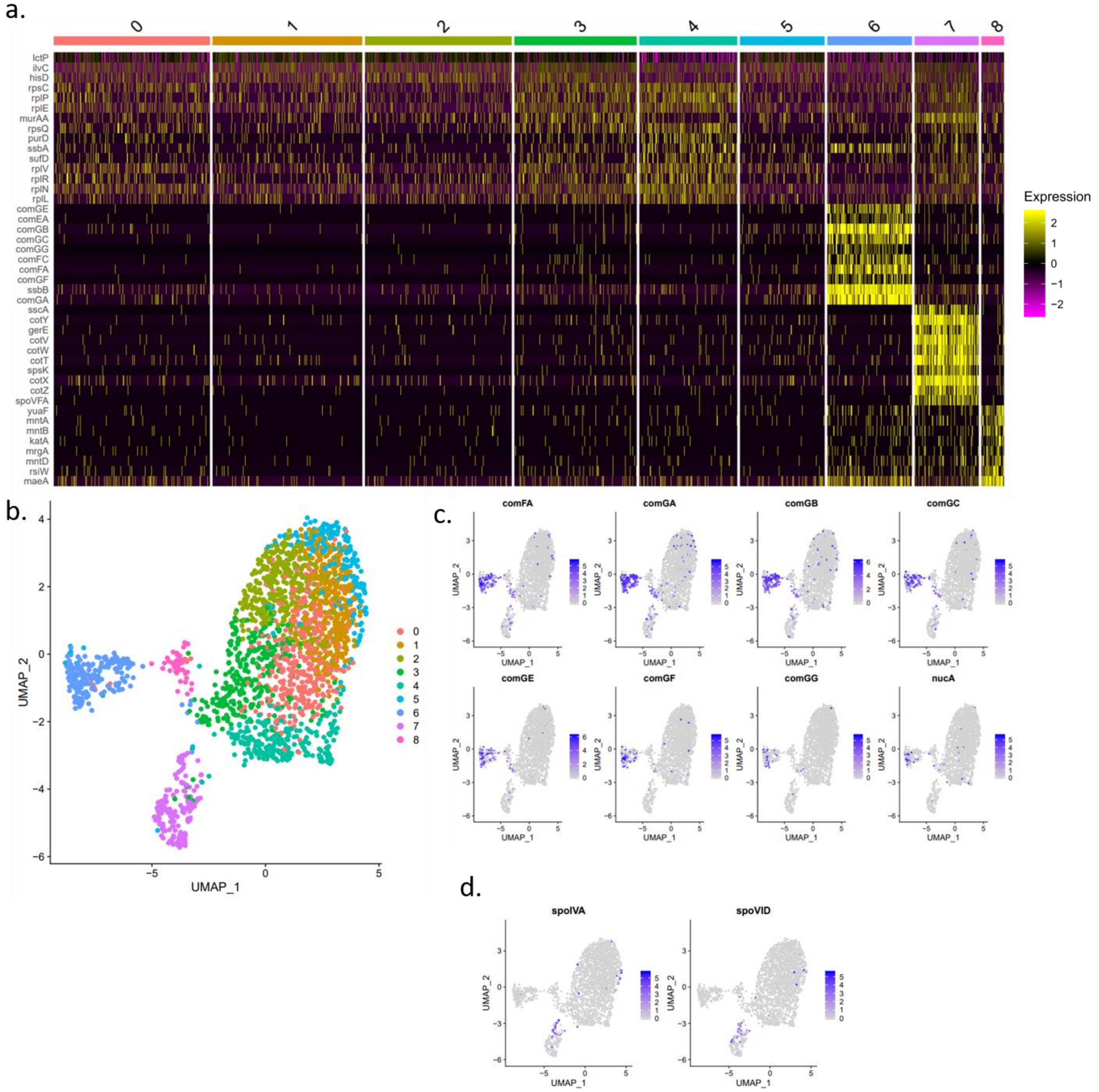


**a.** Heatmap of single cell gene expression **b.** UMAP projection of the 8 clusters **c.** Competence associated genes are predominantly expressed by cells in cluster 6. **d.** Sporulation associated genes are predominantly expressed by cells in cluster 7

**Figure S4:** Heatmap of aerobic M9 culture of E coli cells from figure 4 including all marker gene names

**a.**

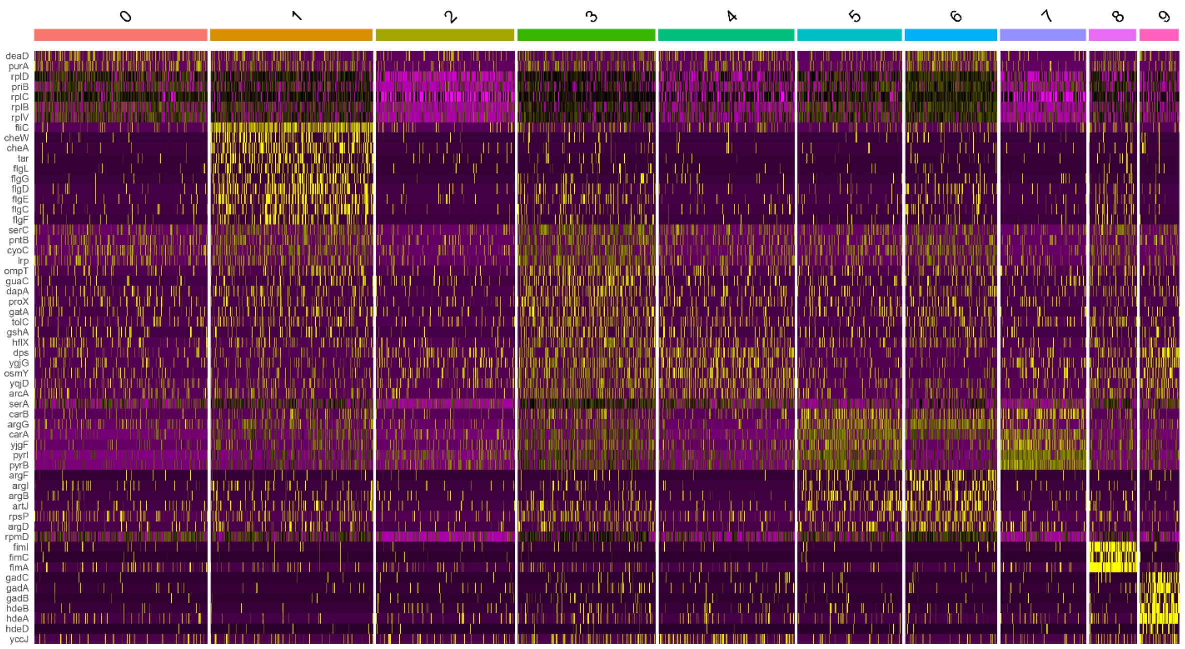


**b.**

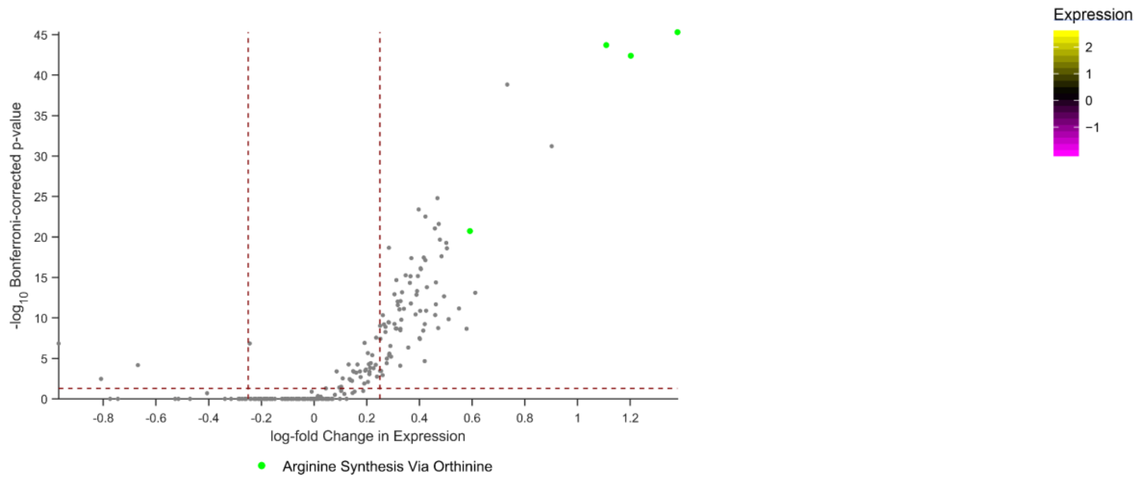


**c.**

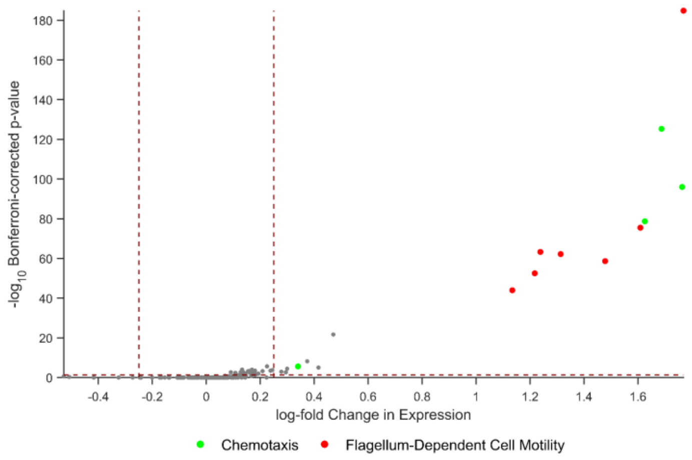


**a.** Heatmap as in figure 4a. including names for all marker genes
**b.** Volcano plot showing gene expression in cluster 6 is enriched for arginine synthesis genes.
**c.** Volcano plot showing gene expression in cluster 1 is enriched for flagella and chemotaxis.

**Figure S5:** Heatmap of E coli fermenting in LB media from figure 4 including all marker gene names

a.

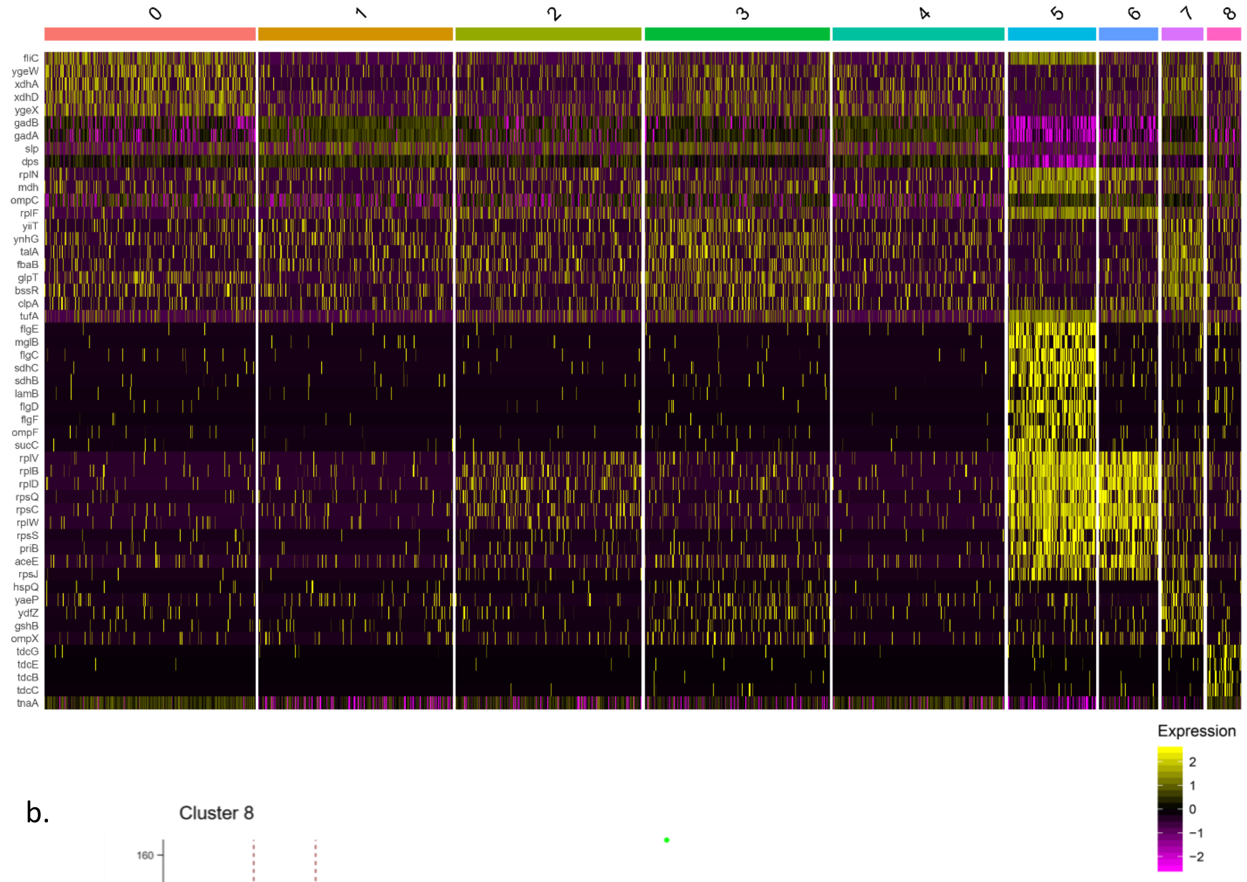


b.

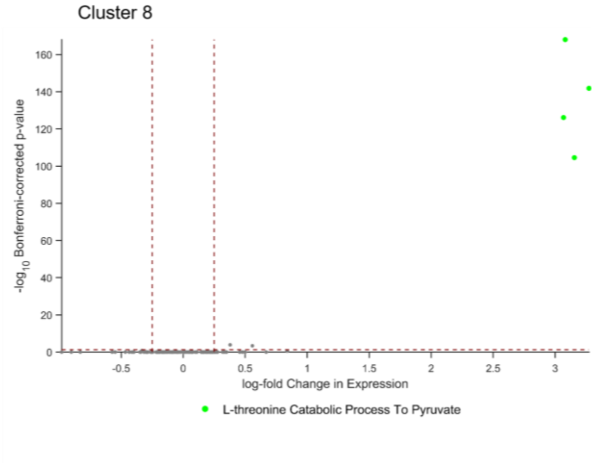


**a.** Heatmap as in figure 4b including names for all marker genes
**b.** Volcano plot showing gene expression in cluster 8 is enriched for L-threonine degradation.

Figure S6: scRNAseq of B subtilis using 10x UMIs instead of Probe UMIs

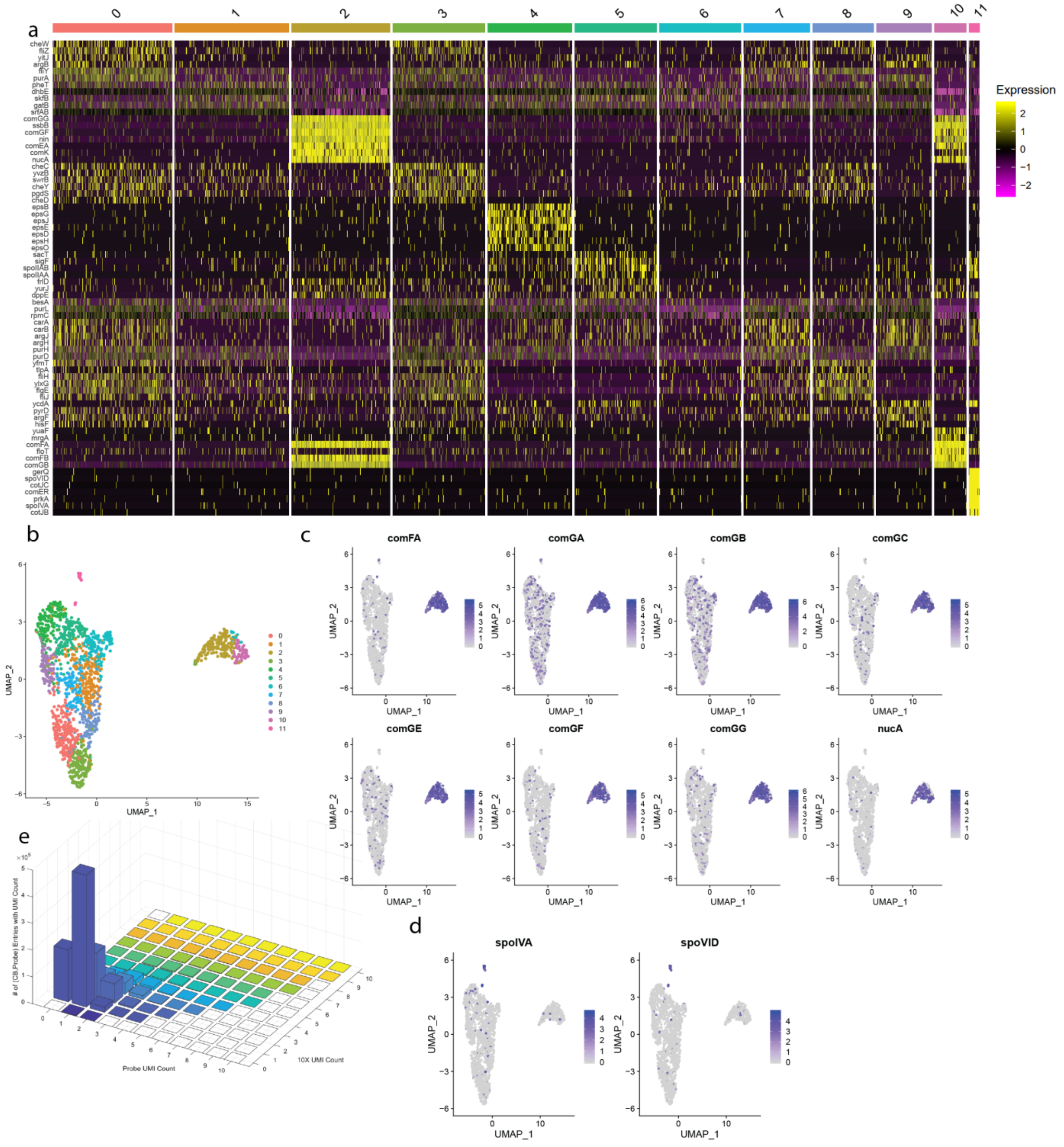


**a.** Heatmap of single cell gene expression **b.** UMAP projection of the 12 clusters **c.** Competence associated genes are predominantly expressed by cells in cluster 2. **d.** Sporulation associated genes are predominantly expressed by cells in cluster 11. **e.** Coincidence histogram of single-cell probe matrix constituted using Probe UMIs vs 10x UMIs (see Methods Section Comparing Single-Cell Probe Expression Matrices Generated Using 10x vs Probe UMIs).

Figure S7: scRNAseq of E coli cells in aerobic M9 culture using 10x UMIs instead of Probe UMIs

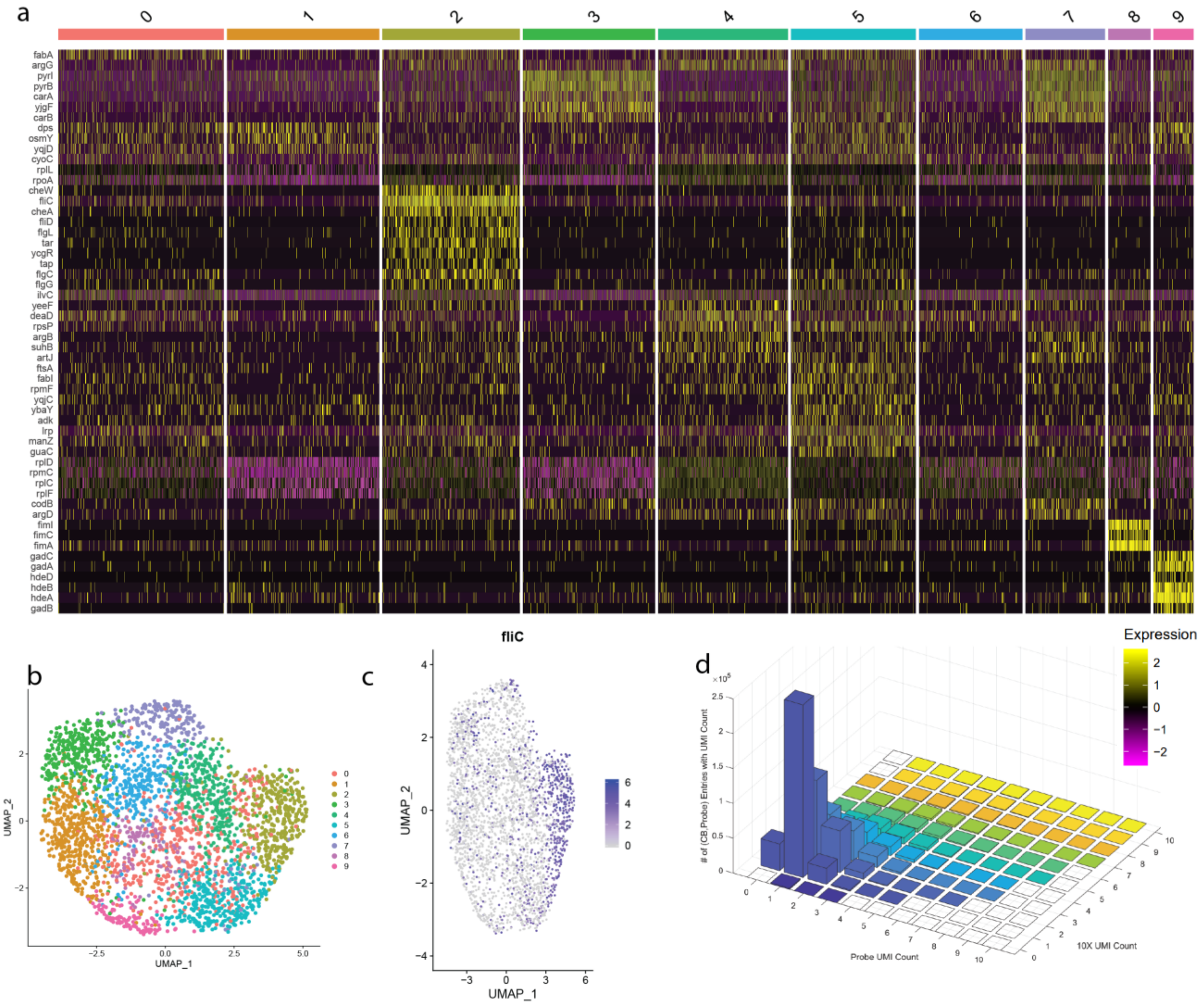


**a.** Heatmap of single cell gene expression **b.** UMAP projection of the 10 clusters **c.** Flagellation associated genes are predominantly expressed by cells in cluster 2. **d.** Coincidence histogram of single-cell probe matrix constituted using Probe UMIs vs 10x UMIs (see Methods Section <u>Comparing Single-Cell Probe Expression Matrices Generated Using 10x vs Probe UMIs</u>).

Figure S8: scRNAseq of E coli cells in LB media using 10x UMIs instead of Probe UMIs

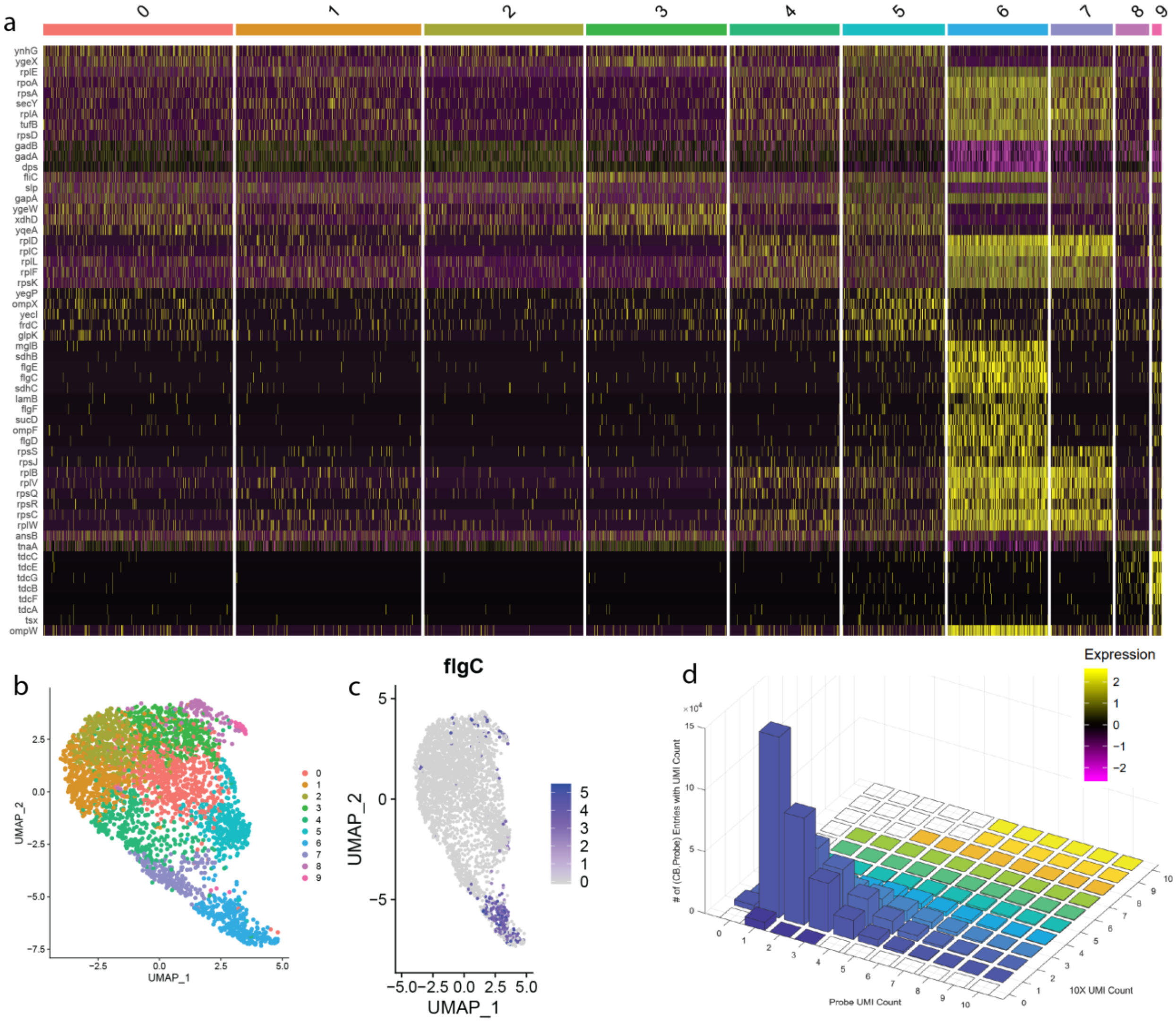


**a.** Heatmap of single cell gene expression **b.** UMAP projection of the 10 clusters **c.** Flagellation associated genes are predominantly expressed by cells in cluster 6. **d.** Coincidence histogram of single-cell probe matrix constituted using Probe UMIs vs 10x UMIs (see Methods Section Comparing Single-Cell Probe Expression Matrices Generated Using 10x vs Probe UMIs).

Figure S9: scRNAseq of B subtilis using bulk median instead of per-cell maximum probe counts for gene expression

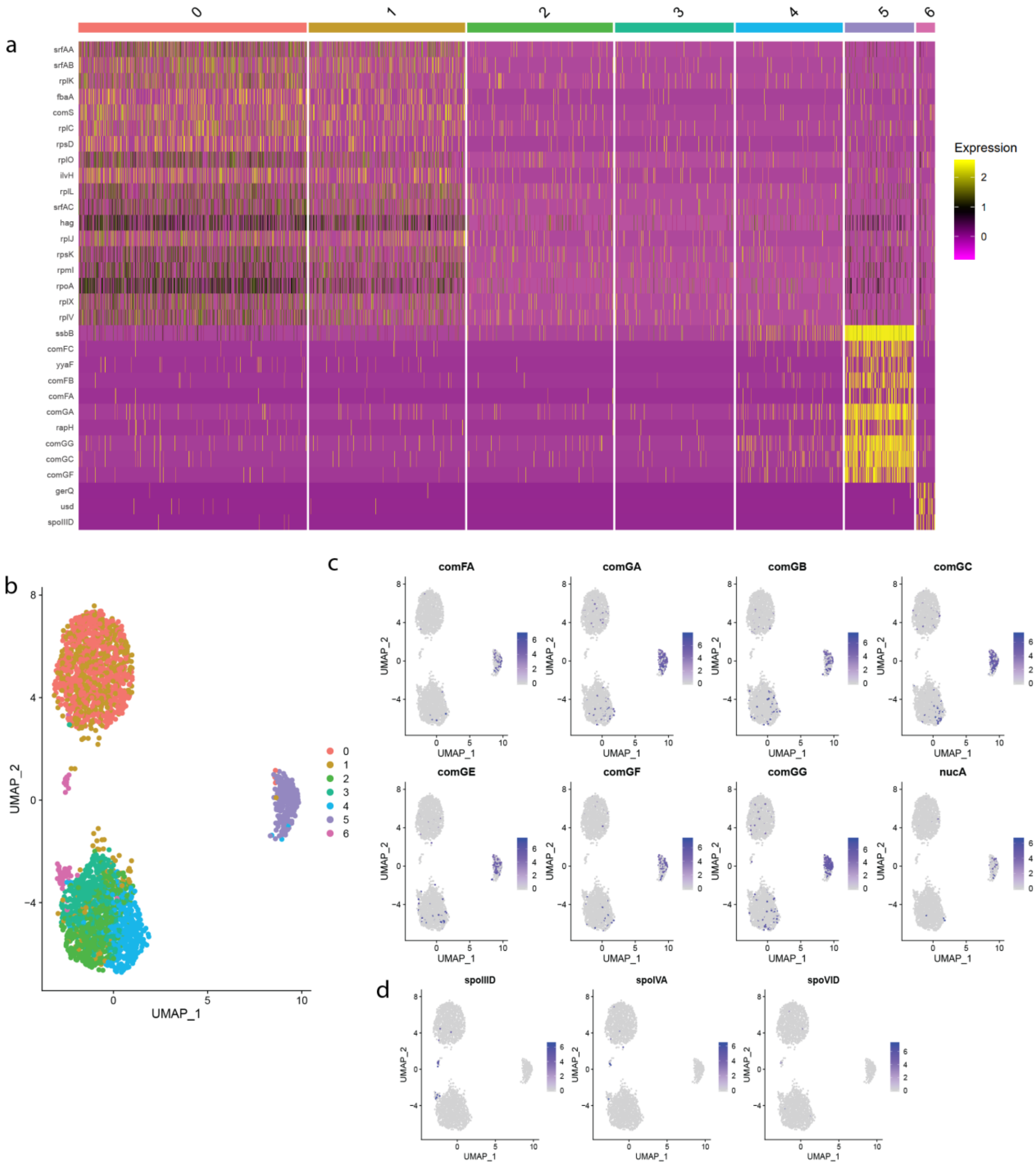


**a.** Heatmap of single cell gene expression **b.** UMAP projection of the 7 clusters **c.** Competence associated genes are predominantly expressed by cells in cluster 5. **d.** Sporulation associated genes are predominantly expressed by cells in cluster 6.

Figure S10: scRNAseq of E coli cells in aerobic M9 culture using bulk median instead of per-cell maximum probe counts for gene expression

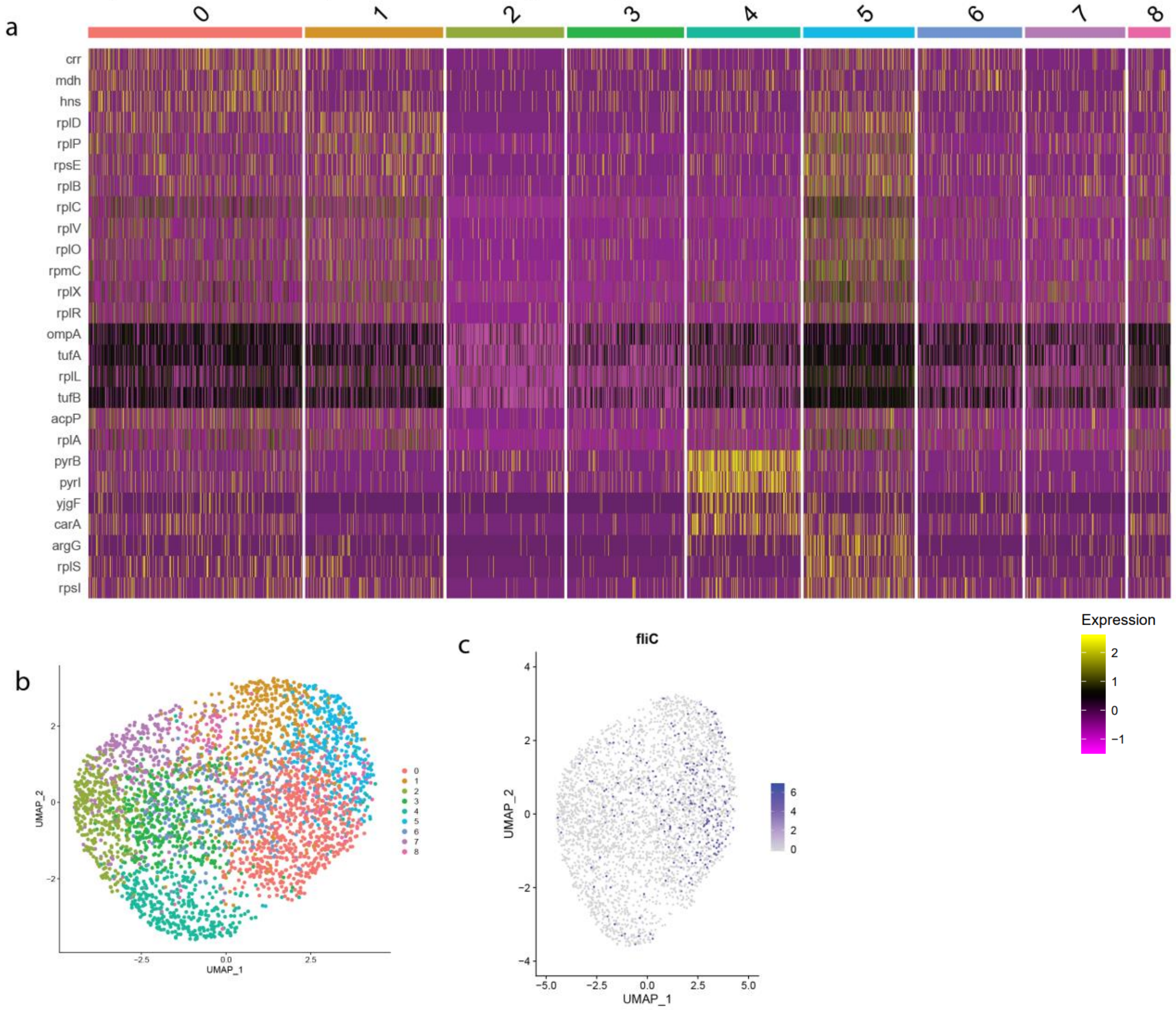


**a.** Heatmap of single cell gene expression **b.** UMAP projection of the 9 clusters **c.** Flagellation associated genes are predominantly expressed by cells in cluster 5.

Figure S11: scRNAseq of E coli cells in LB media using bulk median instead of per-cell maximum probe counts for gene expression

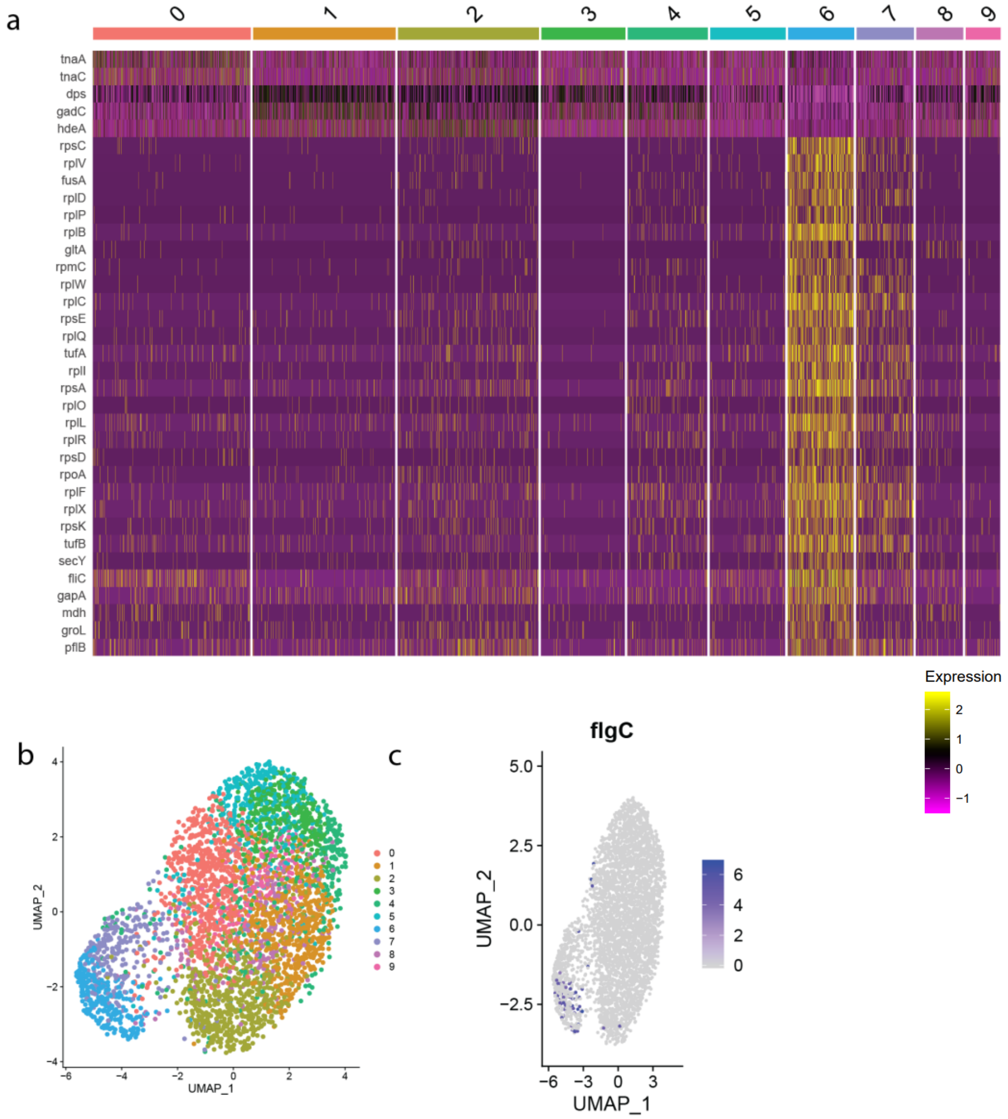


**a.** Heatmap of single cell gene expression **b.** UMAP projection of the 10 clusters **c.** Flagellation associated genes are predominantly expressed by cells in cluster 6.

**Figure S12:** Frequency of marker genes and spores in the population measured by reporter strains

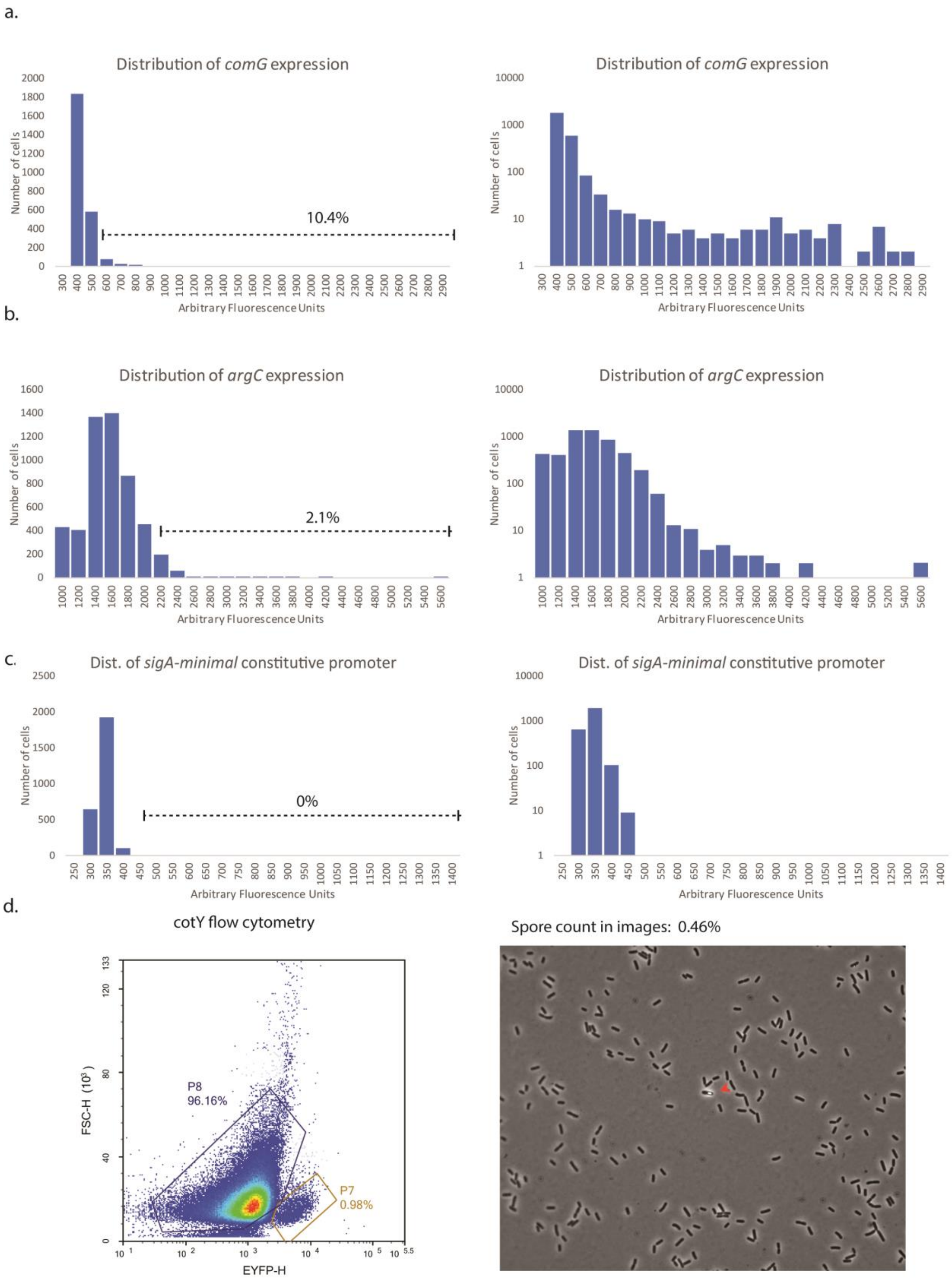

**a.** Approximately 10.4% of the population is in the high-expressing *comG* tail of the distribution (*comG* promoter-reporter strain fluorescence per cell – determined by 1.5x IQR – see Methods). In order to show small cell numbers in the tail more clearly, the histogram is graphed using both a standard scale (left) and a $\log_{10}$ scale (right) for the number of cells in each bin. This is the same strain used in previous studies (Rosenthal et al eLife) with similar distributions **b.** Approximately 2.1% of the population is in the high-expressing *argC* tail of the distribution (1.5x IQR). In order to show small cell-numbers in the tail more clearly, the histogram is graphed using both a standard scale and a $\log_{10}$ scale **c.** A minimal constitutive promoter reporter (sigA) does not have a long-tailed distribution. To show the lack of a long tail the X axis was extended **d.** Sporulation marker gene *cotY* is expressed in approximately 1% of cells (P7 depicts fluorescence from this promoter-reporter strain in the flow cytometry panel, left) and spores are present in a small fraction of cells (approx. 0.5% - right panel)
